# Consistent population activity on the scale of minutes in the mouse hippocampus

**DOI:** 10.1101/2021.02.07.430172

**Authors:** Yue Liu, Samuel Levy, William Mau, Nitzan Geva, Alon Rubin, Yaniv Ziv, Michael E. Hasselmo, Marc W. Howard

**Author notes:** Center for Neural Science, New York University.

## Abstract

Neurons in the hippocampus fire in consistent sequence over the timescale of seconds during the delay period of some memory experiments. For longer timescales, firing of hippocampal neurons also changes slowly over minutes within experimental sessions. It was thought that these slow dynamics are caused by stochastic drift or a continuous change in the representation of the episode, rather than consistent sequences unfolding over minutes. This paper studies the consistency of contextual drift in three chronic calcium imaging recordings from the hippocampus CA1 region in mice. Computational measures of consistency show reliable sequences within experimental trials at the scale of seconds as one would expect from time cells or place cells during the trial, as well as across experimental trials on the scale of minutes within a recording session. Consistent sequences in the hippocampus are observed over a wide range of time scales, from seconds to minutes. Hippocampal activity could reflect a scale-invariant spatiotemporal context as suggested by theories of memory from cognitive psychology.

---

When we remember a particular experience from a trip, other memories from the same trip would also come into mind. Indeed, the retrieval of an episodic memory is believed to involve recovery of the spatiotemporal context associated with that particular episode (Tulving, 1983). The hippocampus has long been implicated in episodic memory (Scoville & Milner, 1957) and it contains single neurons that are active when the animal is at a particular location within an environment (O’Keefe & Dostrovsky, 1971; Moser, Kropff, & Moser, 2008) or at a particular time point during the gap between two stimuli (Pastalkova, Itskov, Amarasingham, & Buzsaki, 2008; MacDonald, Lepage, Eden, & Eichenbaum, 2010; Kraus, Robinson, White, Eichenbaum, & Hasselmo, 2013) (Figure 1a, top). Taken together, this neural population activity can be thought of as a state of spatiotemporal context upon which memories are organized (O’Keefe & Nadel, 1978; Howard, Fotedar, Datey, & Hasselmo, 2005; Polyn & Kahana, 2008; Staresina & Davachi, 2009; Hasselmo, 2012; Eichenbaum, 2017; DuBrow, Rouhani, Niv, & Norman, 2017; Buzsáki & Tingley, 2018; Ekstrom & Ranganath, 2018; Yonelinas, Ranganath, Ekstrom, & Wiltgen, 2019).

**Figure 1.**
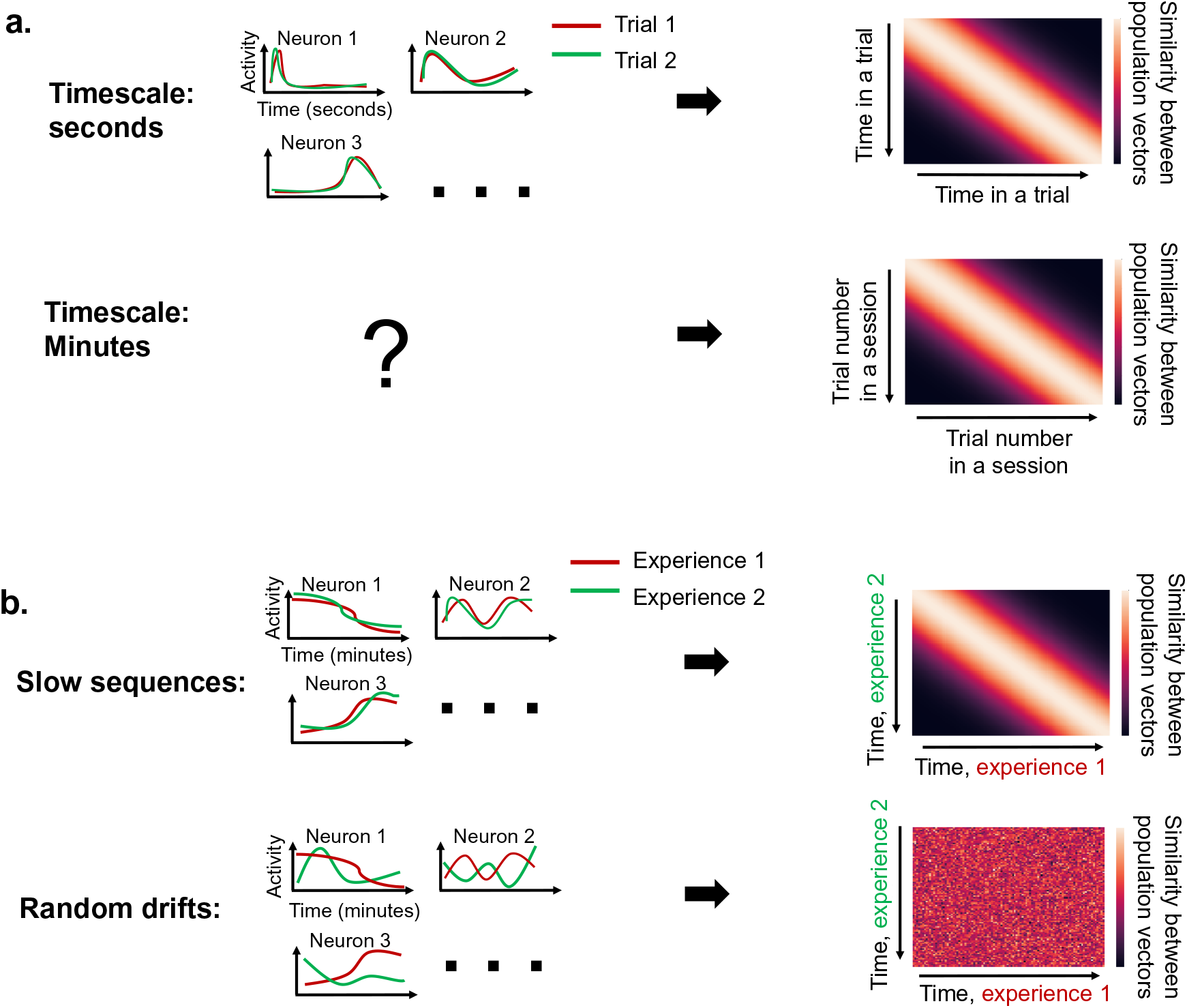
Distinguishing slow consistent sequences from random drifts. **a**. Top: The firing of time cells changes across seconds in sequences that are consistent across trials (left), which contributes to the decorrelation of population activity pattern over the timescale of seconds (right). Bottom: The firing of hippocampal neurons also changes slowly over trials (right), but it is not known if this is driven by consistent sequences on the timescale of minutes (left). **b**. Two possibilities for the nature of the slow dynamics over minutes. Top: it may reflect the animal’s experience. If so, the neural activity would be similar if the animal goes through the same experience twice (left), analogous to the sequences on the timescale of seconds. In this case, the correlation between a pair of population vectors from different experiences will decay as the difference in their time, each within its experience, increases (right). Bottom: alternatively, the slow dynamics may be solely driven by the stochastic noise in neural systems and therefore drift randomly during different experiences (left). In this case, the correlations described above would not have any pattern (right).

Episodic memory retrieval is organized according to spatiotemporal proximity at many different scales. When a participant has an episodic memory for an event from a particular temporal position within a list (Kahana, 1996) or spatial position within an environment (Miller, Lazarus, Polyn, & Kahana, 2013), this brings to mind events from nearby positions, in time or in space. If episodic memory is indeed associated with the recovery of a spatiotemporal context (Tulving, 1983), then the effect of proximity on behavior could be caused by gradual changes in spatiotemporal context reflected in hippocampal ensembles (Manns, Howard, & Eichenbaum, 2007; Ezzyat & Davachi, 2014; Rubin, Geva, Sheintuch, & Ziv, 2015; Cai et al., 2016). In this view, memories for events close in space or time are linked because of overlap in the spatiotemporal contexts associated with those events. Sequences of time cells or place cells could serve as a spatiotemporal context; because they change slowly over time they could mediate associations between nearby events (Wallenstein, Eichenbaum, & Hasselmo, 1998; Howard et al., 2005; Hsieh, Gruber, Jenkins, & Ranganath, 2014). Thus far, time cell sequences have only been observed over a few seconds within the delay period of an experimental trial embedded in a much longer recording session (Pastalkova et al., 2008; MacDonald, Lepage, Eden, & Eichenbaum, 2011; Kraus et al., 2013). However, behavioral effects linking memories separated in time are observed over many timescales in list learning experiments (Howard, Youker, & Venkatadass, 2008; Unsworth, 2008) and can span days and weeks in memory for real-world events (Healey, Long, & Kahana, 2019; Uitvlugt & Healey, 2019). If memories across lists separated by many minutes can be linked, this suggests that hippocampal sequences should also unfold across trials over the scale of minutes. Perhaps the sequence of cells that unfolds in the moments following the beginning of a delay period of a few seconds has an analog in a sequence that unfolds over the entire recording session following the beginning of the session.

A series of studies have found that the activity of hippocampal neurons does change slowly over long timescales. For example, it was reported that population neuronal activity in CA1 exhibits gradual changes over multiple trials that span minutes (Manns et al., 2007; Ziv et al., 2013; Mau et al., 2018) (Figure 1a, bottom). It has also been reported that place cell and time cell activity slowly “drift” across hours and days (Ziv et al., 2013; Mankin, Diehl, Sparks, Leutgeb, & Leutgeb, 2015; Mankin et al., 2012; Rubin et al., 2015; Mau et al., 2018; Cai et al., 2016). The observation of these slow changes with multiple recording techniques make it unlikely that they are a recording artifact. However, it is possible that this slow drift is simply caused by stochastic processes in the neural system or perhaps a gradual but continuous change in the representation of events. Stochastic mechanisms would cause changes in firing across trials but there is no reason to expect that they would cause the same sequence over repeated experiences (Figure 1b, bottom left). However, if slow changes are generated by the same mechanism as time cell sequences, we would expect the dynamics to be consistent across repeated experiences. In much the same way as time cell sequences can be understood as coding for the time since the delay period began, slow changes in firing across trials could contain information about the time since the recording session began.

A more recent study shows evidence for such coding of progression within a session (Sun, Yang, Martin, & Tonegawa, 2020). In that study, mice were trained to run four consecutive laps to obtain a reward. Some neurons in the hippocampus CA1 show elevated activity during a particular lap, and this firing pattern is consistent across repetitions of the same task (Sun et al., 2020). However, it still remains unclear whether similar neural activity patterns can be observed in tasks without a demand to maintain the task progression. It is also interesting to examine whether other forms of temporal modulations are present in encoding the progression of the task. As will be shown in the rest of the manuscript, the answers are positive to both questions.

Figure 1 describes a strategy for data analyses to distinguish consistent sequences from stochastic drifts. Consider a population of cells being recorded over two separate experiences. During each experience the activity of the population changes gradually. This effect can be demonstrated by measuring the correlation of the population activity patterns at different points in time. As one chooses time points further apart from one another within the experience the population becomes more decorrelated. Now, suppose that the population fires consistently from one experience to the next (Figure 1b, top left). In this case one would observe an analogous decorrelation when examining firing across *different* experiences (Figure 1b, top right). Although the two experiences could be separated by a time interval much longer than the duration of the experience itself, the population activity from a particular time point in each experience will be similar even if those time points are taken from different experiences. In contrast, if the within-experience correlation were not due to consistent sequences but, say, stochastic variability (Figure 1b, bottom left), the population would still change gradually over time *within* one experience. However, if the population activity simply decorrelates with time, that would also mean that one would not observe correlation between analogous time points within *different* experiences (Figure 1b, bottom right).

The strategy of this paper is to evaluate whether slow changes in hippocampal activity across trials include consistent sequences extending across multiple trials. In this case, the experience in the example above would be one experimental session that consists of multiple trials. To establish the consistency of the population activity on the scale of a session, it would be necessary to compare the activity between different sessions. By the same reasoning in the example above, if the population activity from a particular time point in each session is similar even if those time points are taken from different sessions, it would indicate that the population firing is consistent between different sessions. Therefore, it requires analysis across multiple sessions across days to analyze the consistency of the neuronal dynamics over minute-level time scales. To compare to the more well-understood sequences—time cells and place cells—we apply the same analyses to the population activity within trials as well. By the same reasoning, it requires analysis across multiple sessions over tens of minutes to examine the consistency of the neuronal dynamics across seconds-level time scales. It is impossible to assess whether a sequence is consistent or not if one cannot record from the same population. Therefore, we study populations recorded using the chronic calcium imaging technique that allows identification of the same neurons across recording sessions (Ziv et al., 2013). We found that across three behavioral tasks, populations of neurons in the CA1 region of mouse hippocampus exhibit consistent dynamics both within second-long trials and across trials, spanning many minutes within a recording session.

## Results

To distinguish consistent sequences from stochastic dynamics (Figure 1), we examined the consistency of the neuronal dynamics across two timescales while mice performed reward-based navigational tasks (Mau et al., 2018; Levy, Kinsky, Mau, Sullivan, & Hasselmo, 2019; Rubin et al., 2015). In all experiments, each session consists of multiple trials, during which mice were trained to navigate through an environment to obtain rewards (Figure 2). One-photon endoscope calcium imaging was used to record the activity of many neurons in the CA1 region of the hippocampus across multiple sessions that span days. Consecutive sessions are separated by at least one calendar day (see Methods section for details). Images from different sessions were cross-registered so that the activity of the same neurons could be tracked across sessions (Methods). We develop a series of computational measures for the consistency of activity over seconds-long delay intervals across trials. Not surprisingly these measures of consistency detect sequences over seconds that are consistent across trials, driven by time cells and place cells. Next, we apply the same computational measures to detect slow sequences of activity over multiple trials that are consistent across experimental sessions. To the extent these measures show the same kind of consistent dynamics described by time cells and place cells, we will establish that hippocampal ensembles exhibit consistent sequences of activity across trials at the time scale of a session.

**Figure 2.**
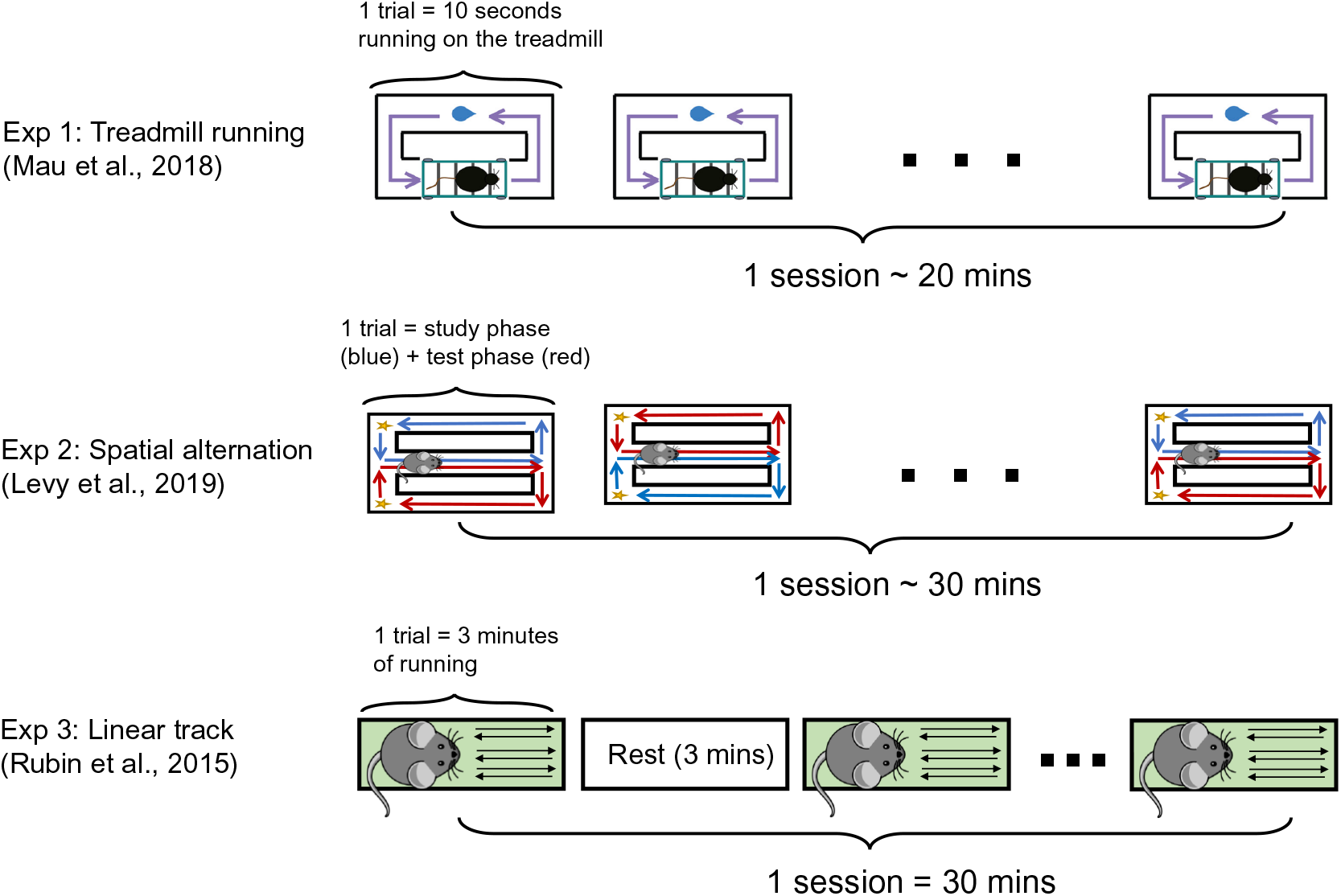
Two timescales in the structure of the experiments. For each experiment studied in this paper, the animals are trained to perform some task for several seconds-long trials in a recording session spanning tens of minutes. The calcium activity of the same neurons are recorded across sessions. During the treadmill running task (Experiment 1), mice are trained to run on the treadmill for 10 seconds before going to the opposite side of the maze to collect a water reward. The mice perform the same task for tens of trials each session for a total of around 20 minutes. For the spatial alternation task (Experiment 2), mice are trained to alternate between left and right turns in a T-maze to collect food rewards. Each trial consists of a study and test phase where mice have to turn to opposite directions at the choice point. Mice perform tens of trials for a total of around 30 minutes during each session. For the linear track experiment, mice are trained to run back and forth on a linear track to collect water rewards at both ends of the track. Each trial is about 3 minutes long and is separated by 3-minute resting periods where mice are placed in a separate box. Each session consists of 5 pairs of running and resting trials for a total of 30 minutes. See Methods section for more details of each experiment.

### Single hippocampal neurons have consistent activity across seconds and minutes

We first extracted the region of interest (ROI), that is, a region of the imaging field believed to correspond to a particular neuron, from the movie obtained from calcium imaging (see Methods for details). Next, we plotted the normalized calcium transient density of individual ROIs against position or time within a trial. For each ROI we only included trials where it had at least one calcium transient event during the trial period examined. We observed that many ROIs have consistent activity within a trial (Figure 3a-c). For example, some ROIs always have higher activity around a particular time bin (Figure 3a, right) or location bin (Figure 3b-c, right) during each included trial. Other ROIs have higher activity at the start (Figure 3a, left) or end (Figure 3b-c, left) of each active trial. At longer time scales, similar consistent neural activity was observed when activity was plotted against trial number (Figure 3d-f, Methods). For example, some ROIs consistently increase (Figure 3d-e, left) or decrease (Figure 3f, left and 3e, right) their calcium activity across trials within a session. Other ROIs are consistently more active during particular trials within the session (Figure 3f, right).

**Figure 3.**
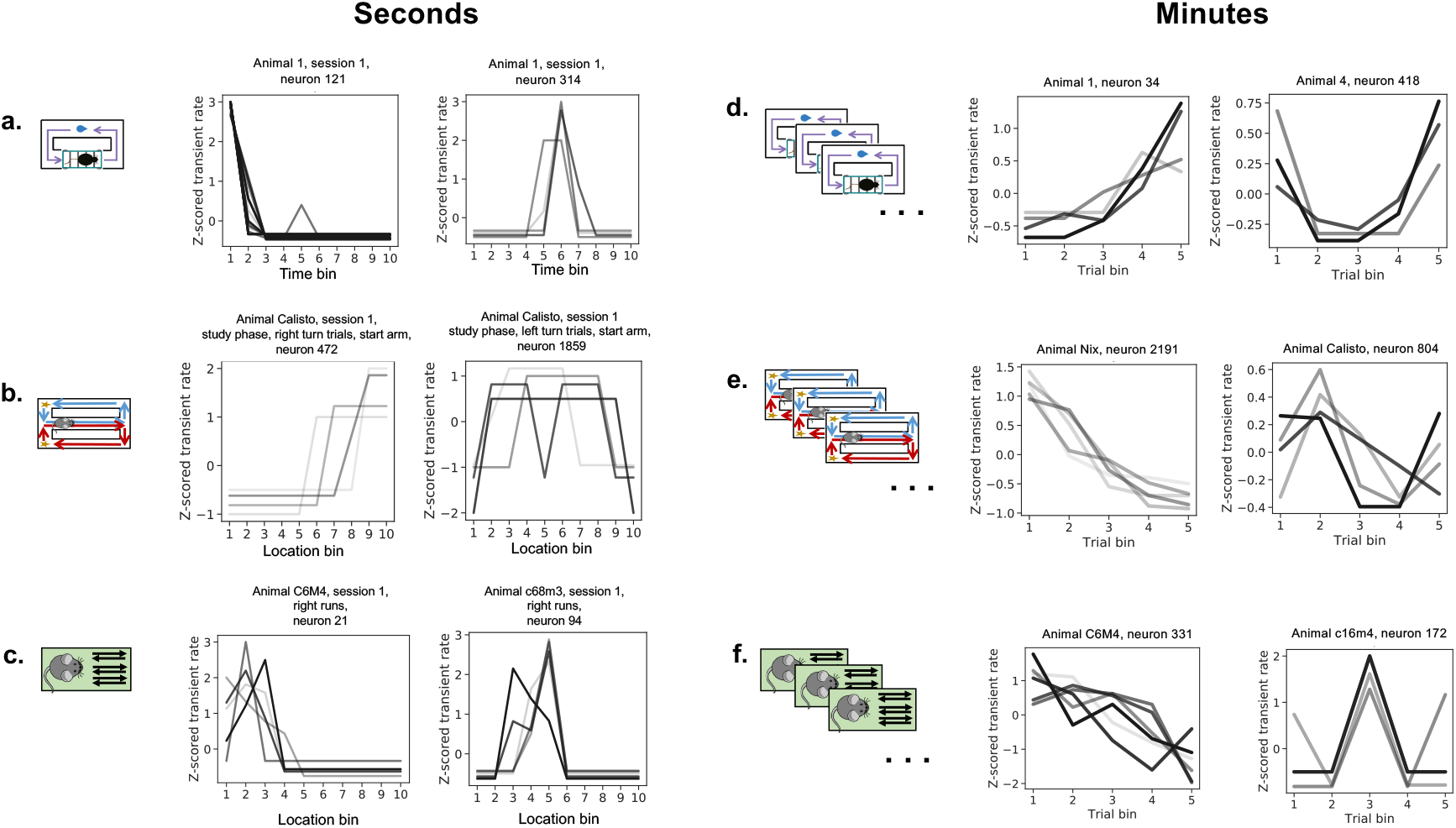
Many hippocampal neurons exhibit consistent dynamics across seconds and across minutes. **a-c**. Example neurons with consistent dynamics across trials for each of the experiments. In **a**, the 10-second running period is evenly divided into 10 time bins. In **b**, the start arm of the maze is evenly divided into 10 location bins. In **c**, the linear track is evenly divided into 10 location bins, and neural activity is averaged over all runs within a 3-minute trial. See Methods for details. **d-f**. Example neurons with consistent activity across sessions. Trials within each session are evenly divided into 5 trial bins. Each line represents the z-scored calcium transient rate over one trial/session. Darker lines indicate earlier trials (for a-c)/sessions (for d-f). Inactive trials/sessions are not shown. See Supplementary figures S1 and S2 for more example neurons.

To quantify the extent to which single neurons fire consistently across the population, we computed a firing consistency score for each neuron, which is a number between zero and one that represents the consistency of that neuron’s calcium dynamics between pairs of trials or sessions compared to chance (see Methods for details). We found that in all experiments and for both timescales, the distributions of the firing consistency score are significantly skewed towards one compared to those obtained from the surrogate data where the mean neural activity during different trial bins was randomly shuffled independently for each ROI (Figure 4). A Kolmogorov-Smirnov test between the distribution from true and shuffled Experie data showed reliable differences in all cases. This test statistic is the maximum distance between the cumulative distribution functions of the two groups of data, and therefore can be regarded as an effect size that is independent of the number of data points. As a comparison, the same statistic between two Gaussian distributions separated by 1 standard deviation is about 0.38. In order to parallel the more frequently used Cohen’s *d* measure for the effect size, we also computed the number of standard deviations *d* that the two Gaussian distributions would need to be separated by for the K-S test statistic to be the same as the ones seen in our data. The results for the above statistical tests are: panel a: *p <* .001, *D*=0.62, *n* = 1860, *d* =1.76; panel b: *p <* .001, *D* =0.61, *n* = 4078, *d* =1.72; panel c: *p <* .001, *D* =0.52, *n* = 1202, *d* =1.42; panel d: *p <* .001, *D* =0.16, *n* = 1338, *d* =0.41, panel e: *p <* .001, *D* =0.13, *n* = 3573, *d* =0.33; panel f: *p <* .001, *d* =0.07, *n* = 1675, *d* =0.18. This indicates there are significantly more neurons that have consistent dynamics both across trials and across sessions than would be expected by chance. The fraction of neurons with a firing consistency score greater or equal to 0.9 among all the neurons included in the analysis is: panel a: 1324*/*1860 = 71.2%; panel b: 2848*/*4078 = 69.8%; panel c: 728*/*1202 = 60.6%; panel d: 324*/*1338 = 25.6%; panel e: 816*/*3573 = 22.8%; panel f: 261*/*1675 = 15.6%.

**Figure 4.**
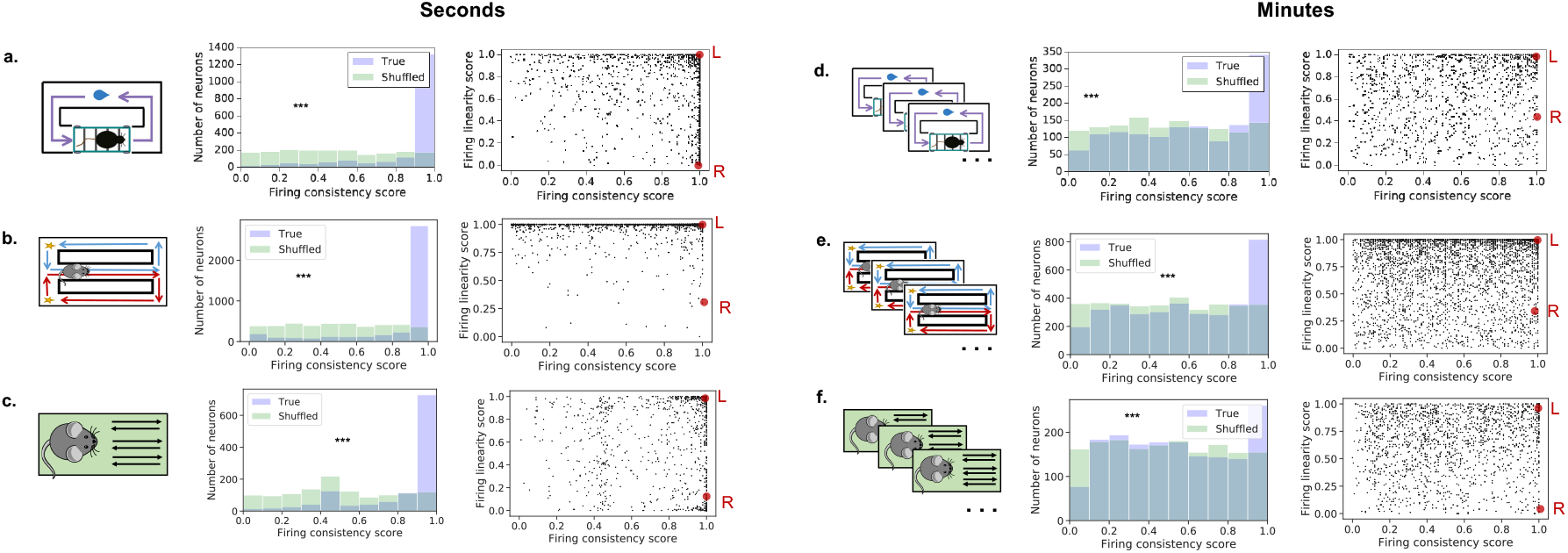
Many hippocampal neurons have consistent responses across seconds and across minutes. For each neuron, a firing consistency score was computed to estimate how consistent the single cell dynamics are for within trial dynamics (**a-c**) and for acrosstrial dynamics (**d-f**). The histograms show the distribution of firing consistency score relative to a surrogate distribution. To the extent these distributions differ, one can conclude that there are consistent sequences. To evaluate the degree to which dynamics were simply monotonic changes in firing rate, we also computed a firing linearity score. The scatter plots show the consistency score and linearity score for each neuron. Red dots indicate the example single neurons shown in Figure 3. L and R refer to the example neuron on the left and right of each panel in Figure 3, respectively. See Methods for more details. Supplementary figures S3-S17 show the same analyses for each trial type, session and animal.

We further confirmed the genuity of this robust long timescale firing by adapting the temporal information metric which was previously used for identifying hippocampal “time cells” on the timescale of seconds (Mau et al., 2018). Briefly, a temporal information score between 0 and 1 was computed based on the Shannon entropy, where 0 indicates that the single cell activity does not reliably carry information about trial number within a session compared to neural activity shuffled across trials (see Methods for details). The distribution of the temporal information score is significantly (*p <* 0.001, Kolmogorov-Smirnov test) different between real and shuffled data in all three datasets (Supplementary figure S20).

There could be different types of single cell dynamics that contribute to the observed high firing consistency across repeated experiences. For example, some cells could have gradually increasing or decreasing activity, or they could exhibit non-monotonic dynamics over time such as those of time cells. To disentangle these two types of single cell dynamics, we used a similar method as above to construct a firing linearity score for each neuron. The right panels of Figure 4 show the joint scatter plot of the firing consistency score and the firing linearity score for each of the experiments. As can be seen, there is a wide distribution of firing linearity across the population. This indicates that there is a diversity of consistent temporal dynamics both across a trial and a session.

We also found that the firing consistency scores on the timescale of minutes and seconds are not significantly correlated with one another across neurons for the linear track task (Supplementary figure S18c, Kendall’s *τ*: *τ* (1099) = −0.01*, p* = 0.6). They are weakly but significantly correlated for the treadmill running and the spatial alternation tasks (Supplementary figure S18a, b. Kendall’s *τ*: *τ* (1245) = 0.13*, p <* 10*^−^*^8^ for a, *τ* (3019) = 0.079*, p <* 10*^−^*^8^ for b). This result implies that the second-scale neurons and the minute-scale neurons are from two independent but overlapping populations in the linear track task, and they show slightly more than chance overlap in the treadmill running and the spatial alternation tasks. In either case, there exist neurons whose dynamics are jointly modulated by two functions of time, one of which changes on the timescale of seconds and one on the timescale of minutes. In other words, there is a multiplexed code for these two different time scales.

### Hippocampal population dynamics are consistent both over seconds and minutes

The single cell analysis above left out trials or sessions where a given neuron is not active (does not have any calcium transient during the selected time period). To examine whether the consistency is present when the full ensemble of neurons are considered, we next investigated the consistency of the population-level dynamics across trials and sessions. To this end, we computed the cosine similarity between pairs of population vectors from different trials (Figure 5a-c) or sessions (Figure 5d-f) and assembled them into a matrix (see Methods for details). We found that in all experiments and across both timescales, the matrices exhibit a pattern where the values are highest along the diagonal, which indicates that the population dynamics are consistent across repeated trials and sessions (c.f., Figure 1b). Statistical significance was evaluated using a permutation test. A “diagonalness score” was computed to quantify the degree to which a matrix shows a diagonal pattern (see Methods for details). As a result, the diagonalness score for all matrices are greater than the matrices obtained from 10,000 random shuffles of the data. We also computed the effect size as the z-scored statistic with respect to the shuffled distribution of diagonalness scores. The values are: Fig 5a: 29.0, Fig 5b: 18.1, Fig 5c: 50.8, Fig 5d: 14.8. Fig 5e: 41.9, Fig 5f: 6.5. Notably, for the across-trial similarity matrix in the treadmill running task, the high-valued region spreads out later in the trial, indicating that the population dynamics slow down as time progresses (Figure 5a). This is consistent with the observation in the original study that the number density of sequentially-activated time cells goes down in time (Mau et al., 2018).

**Figure 5.**
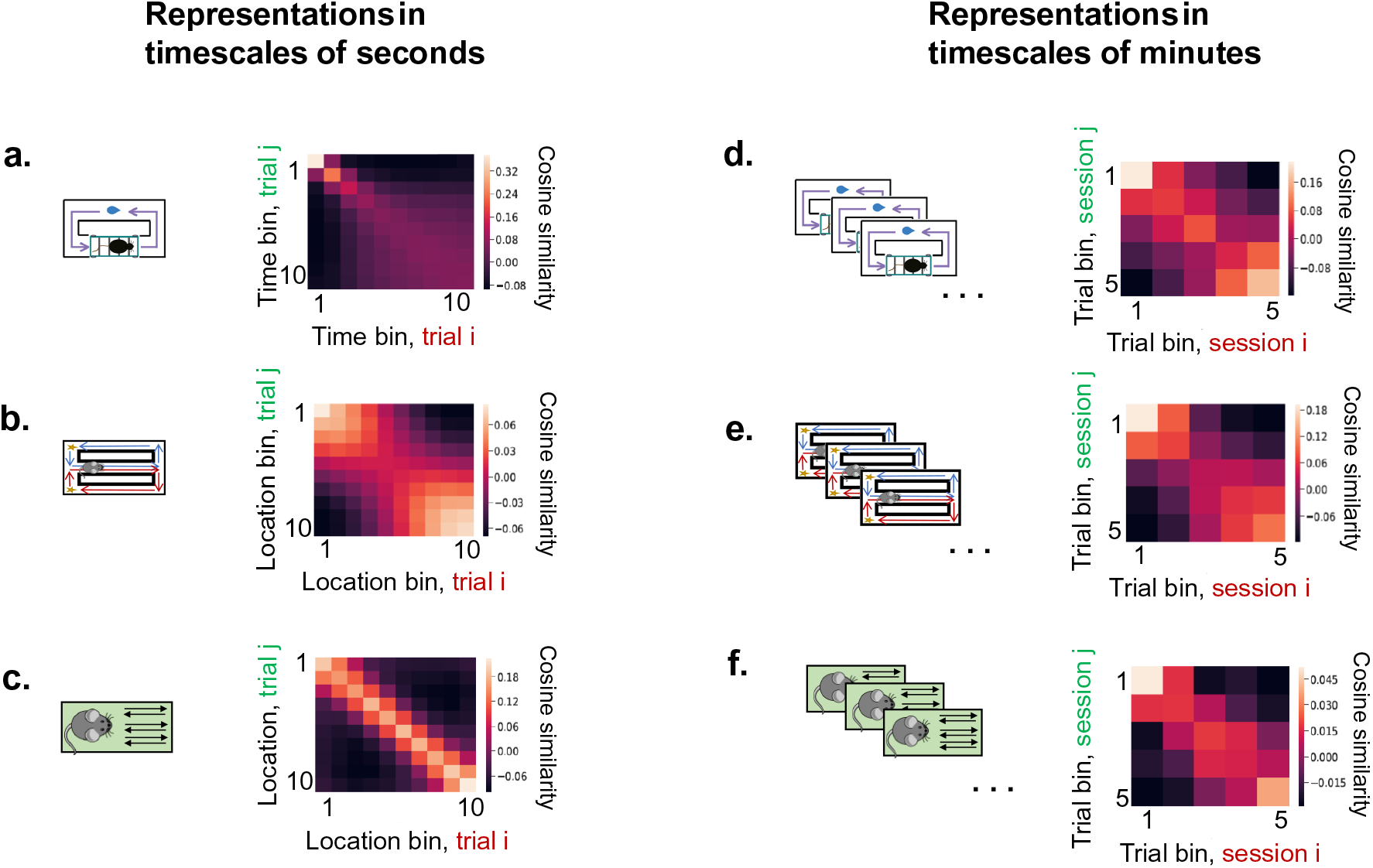
Population dynamics in the hippocampus are consistent across both seconds and minutes. **a-c.** Consistent population dynamics within seconds-long trials. Each element of the matrix is the cosine similarity between a pair of population vectors at different binned locations within a trial, average over pairs of trials, trial types (for b and c), sessions and animals. Critically, the population vectors are taken from the same time or location bin, but from *different* trials. See Methods for the details of how binning was performed for the different tasks. All three experiments (**a**: treadmill task, **b**: spatial alternation, c: linear track) show higher correlation along the diagonal, indicating that the population goes through a consistent sequence within each trial. This is as we would expect from the known properties of time cells (a) and place cells (b and c). **d-f. Consistent population dynamics *across* trials.** Similar to **a-c**, except each element of the matrix is the cosine similarity between a pair of population vectors from two different *sessions*, averaged over all pairs of sessions and animals. Rather than computing population vectors from bins of time or space within a trial, population vectors were computed by averaging over entire trials (see Methods for details). The similarity between population vectors from different recording session was then computed for different pairs of trial bins. The elements of all three matrices are highest on the diagonal and gradually decrease off-diagonal, similar to the matrices over a trial (**a-c**). This indicates that the population dynamics are also consistent across minute-long sessions.

If the population dynamics are consistent across sessions, it should be possible to decode the trial that the animal was in from the population activity. Moreover, this decoder should be generalizable across sessions, meaning that a decoder trained on a subset of sessions should be able to predict the trial number on the rest of the sessions. To verify this, we trained a Linear Discriminant Analysis (LDA) classifier to predict the trial bin number within a session using the neural activity of all the other sessions. The posterior probability given by the LDA classifier for each trial bin was plotted against the actual trial bin (Figure 6, middle). All three heatmaps show significant diagonal patterns. Statistical significance was evaluated using a permutation test similar to the one used for the cosine similarity measure (Figure 5, see Methods for details). The number of shuffles with higher diagonalness scores than the true data is: Fig 6a: 0/1000, Fig 6b: 0/1000, Fig 6c: 5/1000. An effect size was computed based on the z-scored statistic with respect to the shuffled distribution of diagonalness scores, same as for the cosine similarity measure above. The values are: Fig 6a: 4.6, Fig 6b: 11.6, Fig 6c: 2.6. Moreover, the posterior probability given by the LDA for the correct trial bin is above chance for all animals in all tasks (Figure 6, right). These results show that the LDA classifier trained on all but one sessions can accurately predict the trial bin number of the left-out session. In other words, the minute-scale population dynamics over multiple trials are consistent between different sessions. Taken together, both the cosine similarity and the LDA decoding analysis confirm and reinforce the result obtained from the single cell analysis (Figure 4) that the population exhibits consistent dynamics over the timescale of both seconds and minutes.

**Figure 6.**
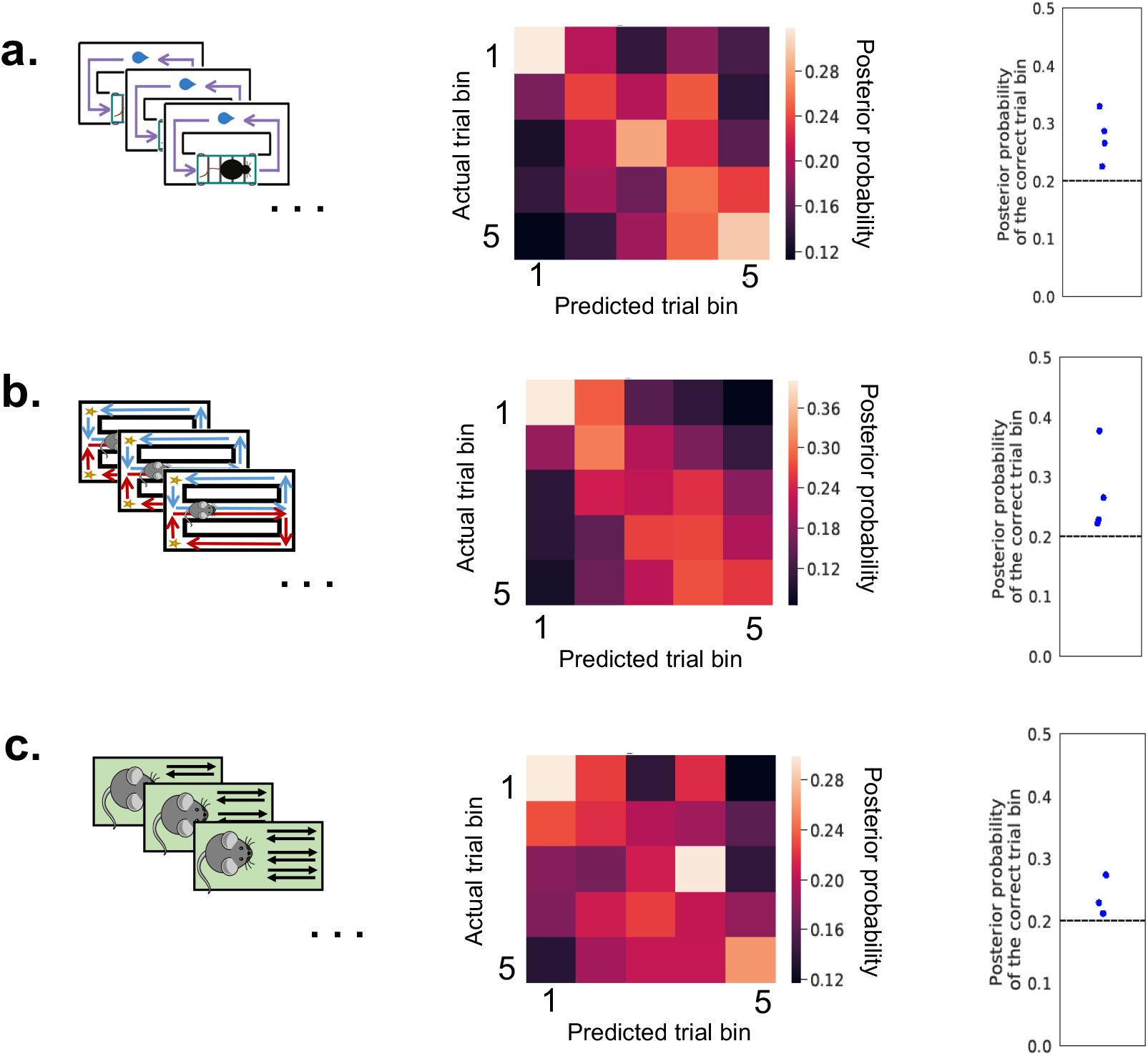
Cross-session decoding for trial bin number. An Linear Discriminant Analysis (LDA) classifier was trained to predict the trial bin number within a given session from the mean population neural activity during that trial bin. The classifier was trained on all but one sessions and tested on the left-out session. Middle: all three heatmaps exhibit a diagonal pattern, indicating that the classifier can correctly predict the trial bin number within the left-out session. The diagonal pattern in the heatmaps indicate successful decoding. Statistical significance was evaluated by computing a diagonalness metric (same as the one used for the heatmap of correlations. See Methods) and comparing with shuffled data. Right: the posterior probability for the correct trial bin given by the LDA classifier (each point represents an animal) compared to chance ( 1/5 = 0.2, dashed line).

## Discussion

In this paper, we show that the firing dynamics of hippocampal neurons are consistent over both seconds and minutes. The novel observation is that slow dynamics over minutes include slow sequences and are not simply random drifts (Figure 5d-f, Figure 1b). This population effect results from a significant proportion of neurons with consistent dynamics over repeated experiences (Figure 4). These neurons have both monotonic and more complex activity modulations across each experience (Figures 3 and 4 and Supplementary figures S1 and S2). Therefore, the hippocampal neurons exhibit consistent dynamics over two nested timescales—changing both systematically within a trial and systematically within a session—in each of these experiments.

As seen above, the effect sizes for the population analysis are much higher than those for the single cell analysis. This is because the effect size for the single cell analysis is more affected by the variability across cells in terms of their firing consistency, whereas the one for the population analysis measures the significance of the firing consistency on the population level, therefore less affected by single cell variability.

Multiple metrics were used to quantify the firing consistency on the single cell and population levels. The single cell rank metric (Figure 4) directly measures the correlation between the temporal firing patterns of two experiences. The temporal information metric (Figure S20) measures the temporal modulation of the average firing across experiences, and is commonly used to identify second-scale time cells (e.g. Mau et al., 2018). The two population analyses also serve slightly different purposes: the cosine similarity metric (Figure 5) directly measures the consistency of the population temporal firing patterns across two experiences. On the other hand, the LDA classification accuracy metric (Figure 6) measures the ability for the same downstream population to linearly read out elapsed time on the scale of seconds across trials, and to read out task epoch on the scale of minutes across sessions.

This result suggests that the spatiotemporal context as represented by population of neurons in the hippocampus has meaningful dynamics over multiple timescales, from seconds to minutes. The sensitivity to multiple timescales may enable the hippocampus to adaptively encode natural stimuli, which vary at many different scales (Voss & Clarke, 1975; Hasson, Yang, Vallines, Heeger, & Rubin, 2008) and account for the self-similar structure in hippocampal correlations (Meshulam, Gauthier, Brody, Tank, & Bialek, 2019). The responsiveness of hippocampal dynamics provides a constraint for behavioral models of human memory. Models that rely on boundaries and event segmentation (Farrell, 2012; Franklin, Norman, Ranganath, Zacks, & Gershman, 2020) must be able to construct and utilize segments over multiple nested scales. Similarly, neural models for sequence generation (Buzsáki & Tingley, 2018; Rajan, Harvey, & Tank, 2016; Howard et al., 2014) must have the capacity to generate sequences at many different scales.

A recent study shows that there are neurons in the hippocampus CA1 of mice that encode the number of laps that the animal has traversed in a task where they have to run four consecutive laps to obtain a reward (Sun et al., 2020). These “lap cells” also constitute a reproducible hippocampal sequence, but our results differ from this study in three aspects. First, while some of the neurons reported in our work indeed show elevated activity for a particular trial bin, analogous to the “lap cells” in Sun et al., the majority of neurons that we observed exhibit other types of temporal modulations (Figure 3d-f and Supplementary figures S1-S2). Second, the information about trial number is beneficial for the mice in the task of Sun et al., yet is not useful for the mice in the tasks we analyzed. Yet, slow dynamics are still observed. Third, the timescale of the slow dynamics in our study is an order of magnitude longer than that observed in Sun et al..

### Possible causes of slow dynamics

#### Possible non-physiological causes

There are recording artifacts specific to calcium imaging that could conceivably cause slow dynamics. For example, photobleaching could cause the calcium fluorescence signal to decrease gradually during each imaging session or gradual heating of the brain could potentially produce stereotypical changes in the apparent calcium fluorescence signal for each ROI over the course of an imaging session. It is difficult to reconcile these artifactual accounts of slow dynamics with non-monotonic patterns of firing over the session or the similarity between effects observed across trials to the effects within trial. The findings within-trial are quite consistent with results using extracellular electrodes. We conclude that it is likely that the slow dynamics are physiological in origin.

#### Variables correlated with time during a session

It is possible that the slow dynamics observed in the hippocampus are not due to memory *per se* but reflect consistent slow dynamics in the environment or internal state of the animal over the course of the recording session. There are several possibilities for such variables. For example, the satiety of the animal presumably decreases over the course of each recording session. Indeed, it has recently been reported that thirst level has a dramatic impact on the population activity in multiple brain regions in mice over the course of minutes (Allen et al., 2019). Moreover, a recent study shows that slow drifts over minutes in area V4 and prefrontal cortex of monkey are correlated with systematic changes in animal’s behavior during a perceptual decision making task (Cowley et al., 2020). In addition, microdialysis of acetylcholine shows higher levels of acetylcholine when an animal is first removed from the home cage and placed in a task (Acquas, Wilson, & Fibiger, 1996). Acetelcholine levels decrease over time on the scale of minutes and have been shown to depolarize hippocampal neurons (Cole & Nicoll, 1984) and increase firing rate (Fu et al., 2014), consistent with the cells that gradually increase or decrease their activity (Figure 3, 4). In all of these cases, the sequential activation of cells over the recording session would require that the hippocampus codes for a monotonically changing variable with a sequence of receptive fields. Indeed, this kind of pattern has been observed for hippocampal receptive fields as a function of smooth changes in frequency of a behaviorally-relevant tone (Aronov, Nevers, & Tank, 2017).

In some sense the empirical story for very slow sequences is analogous to the empirical story for place cells or time cells shortly after their initial report. Although a consensus has emerged that place cells and time cells express spatial and temporal relationships between events in the service of memory (e.g., Eichenbaum, 2017), this view only emerged after extensive empirical studies. For instance, a neuron that fires when the animal is in a specific position of an environment could be responding to the visual stimuli available at that location, the particular configuration of auditory stimuli available, or olfactory cues present on the track. Early studies ruled out a series of possible confounds of spatial position (e.g., Quirk, Muller, & Kubie, 1990; Save, Cressant, Thinus-Blanc, & Poucet, 1998). Similarly, it is possible that initial reports of time cells could have been solely a reaction to a behavioral confound during the delay such as a stereotyped behavior. However, time cells have been observed in head-fixed animals and different stimuli trigger distinct sequences (e.g., Pastalkova et al., 2008; Taxidis et al., 2020; Cruzado, Tiganj, Brincat, Miller, & Howard, 2020), ruling out most possible confounds.

#### Slow dynamics as memory for the past

It has been clearly established that hippocampal time cells express memory for the time and identity of past events (e.g., Pastalkova et al., 2008; Taxidis et al., 2020; Cruzado et al., 2020). The most interesting possible cause of the slow dynamics observed here is that they reflect the same computational mechanism, but over much slower time scales than within-trial time cells. How might the same computational mechanism generate neural dynamics across a wide range of timescales? It has been suggested (Rolls & Mills, 2019; Shankar & Howard, 2012) that sequential activity across multiple timescales could originate from cells that show exponential decaying activity over the same range of timescales, which have been reported in two recent studies. Tsao et al. (2018) observed slow changes in the firing of neurons in lateral entorhinal cortex (LEC). In that study, LEC neurons changed their firing rate abruptly and then relaxed back to baseline with a broad range of decay rates. For instance, upon entry to a particular environment, a neuron might rapidly increase its firing rate and then decay exponentially back to baseline over several minutes. Other neurons ramped over the entire recording session so there was a variety of decay rates across neurons. This slowly-varying signal in LEC at the scale of minutes could be a cause of the slow sequences we observed in hippocampus. In another study, Bright et al. (2020) studied neurons in monkey EC during a visual task. After an image was presented, the neurons changed firing rate then gradually relaxed back to baseline with a variety of decay rates. Because there was a variety of relaxation rates it was possible to decode time since image onset over several seconds (see also Hyde & Strowbridge, 2012). Very long-lasting firing in EC has been observed *in vitro* (Egorov, Hamam, Fransén, Hasselmo, & Alonso, 2002) and is believed to be caused by the nonspecific calcium-sensitive (CAN) cationic current. Computational models show that the CAN current can also induce slowly decaying firing with a variety of decay rates in a simple integrate-and-fire neuron model (Tiganj, Hasselmo, & Howard, 2015). Computational models have shown that the temporal information carried by slowly-decaying activity can be used to generate a population of sequentially-activated time cells (Shankar & Howard, 2012; Howard et al., 2014; Liu, Tiganj, Hasselmo, & Howard, 2019; Rolls & Mills, 2019; Liu & Howard, 2020). Of course, the definitive test of whether the slow hippocampal sequences reflect a very slow form of memory is whether the identity of events on previous trials can be decoded. This analysis is not feasible given the design of the tasks analyzed here, but similar analysis have been done on the neural activity from other cortical regions (e.g. Schoenbaum & Eichenbaum, 1995b, 1995a; Bernacchia, Seo, Lee, & Wang, 2011; Morcos & Harvey, 2016).

In summary, it was shown that the slow neuronal activity on the timescale of minutes are consistent across repeated sessions. This slow dynamics is most likely related to either a gradual change of the animal’s internal state, or a gradual evolution of the animal’s memory trace about past events, or both. The exact function of and the mechanism that generates this slow consistent activity remains to be elucidated in future experiments.

### Methods Behavioral tasks and calcium imaging

#### The treadmill running task

All procedures were in compliance with the guidelines of the Boston University Animal Care and Use Committee. Four mice were trained to traverse a rectangular track followed by running in place on a motorized treadmill for 10 s at a constant velocity to receive sucrose water reward after traversing an additional part of the track (Figure 2, Experiment 1). During each session, the mice completed between 35-37 trials. A total of 4 sessions were performed for each mouse.

Mice received infusions of AAV9-Syn-GCaMP6f (U Penn Vector Core). Imaging data in dorsal CA1 were acquired using a commercially available miniaturized head-mounted epifluorescence microscope (Inscopix). The raw video was pre-processed using an image segmentation algorithm called Tenaspis (D.W. Sullivan et al., 2017, Soc. Neurosci., abstract, software available at https://github.com/SharpWave/TENASPIS) to extract ROIs and assign calcium transient events to each ROI. This algorithm is designed to better distinguish between overlapping ROIs. The calcium transients it detects correspond to the rising phase of the calcium fluorescence. 296-1136 ROIs were identified during each session. There is a total of 4 imaging sessions spanning 4 calendar days.

In order to identify the same neurons across recording sessions that are days apart, ROIs were cross-registered across days. Briefly, this was done by first aligning the field of view of each session to the first session using vasculature as stationary landmarks via image registration software from MATLAB’s Image Processing Toolbox, assuming rigid geometric transformation. Then, cells were successively registered from each session to the next session (Day 1 to Day 2, Day 2 to Day 3, etc.). Cells were registered by searching for the nearest ROI with a threshold that the ROI centroids must be within 3.3 microns apart. To ensure that neurons do not drift excessively across days, the first day’s neurons were registered with the last day’s neurons, and any registrations between Day 4 and Day 1 that are different from Day 4 and Day 3 were discarded. Cells are stably tracked across sessions using this method, as illustrated in the original paper (Figure 4, 5 and S3 in Mau et al., 2018). More details about the behavioral setup and the calcium imaging experiment can be found in the Methods section of Mau et al. (2018).

#### The spatial alternation task

All procedures presented here were approved by the Institutional Animal Care and Use Committee (IACUC) at Boston University. Four mice received infusions of AAV9-Stn134 GCaMP6f (University of Pennsylvania Vector Core, obtained at a titer of 4 10^13^GC/mL and 135 diluted it to 5 6 10^12^GC/mL with 0.05 M phosphate buffered saline). They were trained on a spatial alternation task, during which they alternated between “study” and “test” trials. On study trials, mice were placed on the center stem of maze, ran to the crossroads, where a removable barrier forced them to run down one of the two return arms and received a reward of chocolate sprinkle. They were then moved into the delay area located at the bottom of the center stem, waited through a 20-second delay, and the delay barrier was lifted to start the test trial. On a test trial, mice again ran up the center stem to the crossroads but this time there was no barrier and they had to remember the direction they traveled on the study trial and turn to the return arm opposite to the preceding study trial in order to receive a reward. They then moved to the delay area, and were placed in their home cage to wait through a 15-25 second inter-trial interval while the next study trial was set up (Figure 2, Experiment 2). Mice completed between 25 and 40 study-test trial pairs per session. Each of these trial pairs is considered a “trial” in the analysis in the main text. There is a total of 9-11 imaging sessions spanning up to 17 calendar days

The experimental procedures for calcium imaging and the data pre-processing pipeline are the same as the treadmill running task. 1149-3165 ROIs were identified for each session. The cell cross-registration procedure is slightly different from the treadmill running task. Sessions were aligned to a “base” session from the middle of the recording schedule using 25-40 “anchor” cells. Cells with centers within 3 microns were identified as the same cell. Cells are stably tracked across sessions using this method, as illustrated in the original paper (Figure 1 and Supplemenatary Figure 1 in Levy et al., 2019). More details about the experimental setup can be found in Levy et al. (2019).

#### The linear track task

All procedures were approved by the Weizmann Institute IACUC. Three mice (2 were injected with AAV2/5-CaMKIIa-GCaMP6f and one was a Thy1-GCaMP6f transgenic; Jackson stock number 025393) were trained to run back and forth on an elevated 96 cm long linear track. They received water sweetened with lemon flavored fruit juice concentrate at each end of the track. An overhead camera (DFK 33G445, The Imaging Source, Germany) was used to record mouse behavior. Each session consisted of five 3-min trials with 3-min intertrial intervals. There are 7-8 imaging sessions conducted every other day for each mouse. Sessions are in the morning and the afternoon in alternation. An integrated miniature fluorescence microscope (nVistaHD, Inscopix) was used to obtain the imaging data from the CA1 region of the hippocampus. Imaging data was preprocessed using commercial software (Mosaic, Inscopix) and custom MATLAB routines as previously described in Ziv et al., 2013. Spatial filters corresponding to individual ROIs were first identified using a cell-sorting algorithm that utilizes principal component analysis and independent component analysis (PCA and ICA, Mukamel, Nimmerjahn, & Schnitzer, 2009) and then subjected to further manual cell sorting (see the “Materials and methods” section of Rubin et al. (2015) for more details). Calcium transient events were identified when the amplitude of the calcium traces 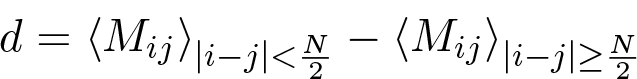 crossed a threshold of 5 median absolute deviations (MAD). Further measures were taken to avoid the detection of multiple peaks as well as the spillover of the calcium fluorescence to neighboring cells. More details about the method can be found in the “Materials and methods” section of Rubin et al. (2015)

Registration of cells across sessions was performed by first aligning the field of view in each session to the first session and then computing the spatial correlation between ROI centroids in the reference coordinate system. Pairs with spatial correlation *>* 0.7 or distance *<* 5 *µm* were registered as the same neuron. Cells can be stably tracked over days, as illulstrated in the original article (Figure 1 and Figure 1-figure supplment 3 in Rubin et al., 2015). For the full detail on the experimental setup please refer to the “Materials and methods” section of Rubin et al., 2015.

## Data analysis methods

### Coarse-graining of calcium activity

#### Coarse-graining for the across session dynamics

To extract the slow neuronal dynamics across multiple trials while filtering out the fast within-trial dynamics, the neural activity was first temporally coarse-grained before further analysis. When comparing a pair of sessions, the session with more trials was first truncated to have the same number of trials as the other session. Then, all the remaining trials within a session were evenly divided into 5 trial bins by using the `array_split` function in `Numpy`. Then the neural activity for each ROI is the calcium transient density averaged over that trial bin. Therefore, the temporally coarse-grained activity of an ROI *n* during a session *i* was represented by a time series with length 5: *v_n,i_* = *v_n,i,_*_1_*, v_n,i,_*_2_*, v_n,i,_*_3_*, v_n,i,_*_4_*, v_n,i,_*_5_ . Furthermore, since we are interested in the temporal modulation of the neural activity rather than the absolute magnitude of the activity, the coarsed-grained activity of each cell was z-scored across trial bins. We chose 5 as the number of time bins within a session since each session in the linear track task consists of 5 running trials (Figure 2, Experiment 3), and we wish to keep the way trials are divided consistent across experiments. Similar results were obtained for the treadmill running task and the spatial alternation task by using 10 trial bins. Furthermore, we only averaged over the calcium activity during time periods when the animal’s behavior is under experimental control. In the treadmill running task, the time periods used are when the animal is running on the treadmill for 10 seconds. In the spatial alternation experiment, the time periods used are when the animal is running along the start arm. In the linear track experiment, the time periods used are when the animal’s position is within the middle 60% of the linear track.

#### Coarse-graining for the within-trial dynamics

To extract the within-trial neuronal dynamics, coarse-graining was done in a similar way by computing the calcium transient density over 10 time bins or location bins within each trial. For the treadmill running task, calcium event rate was averaged over each second during the 10-second running period. For the spatial alternation task, the start arm was evenly divided into 10 location bins and total number of calcium transient events within each bin divided by the amount of time the animal spent in that bin was computed. Unless otherwise specified, all analysis was performed separately for the two task epochs (study and test) and two trial types (turn left and turn right) and the results were averaged. For the linear track task, the within-trial neural activity was computed by first computing for each individual run the number of calcium transient events within each location bin divided by the amount of time the animal spent in that bin, and then averaging this quantity over all runs within a 5-minute trial. This was done for the two running directions separately, and the results were averaged. The 10 location bins span the middle 30% of the track. We chose the middle 30% of the linear track because this is similar to the length of the start arm in the spatial alternation task. Lastly, for all experiments, the activity of each neuron was z-scored across all spatial or time bins for each trial.

#### Population analyses

To quantify the consistency of the population dynamics across sessions (Figure 5d-f), we computed the cosine similarity between pairs of population activity vectors during different sessions after they were coarse-grained and z-scored as described above. We then built a matrix where each element represents the cosine similarity between a pair of population vectors at two trial bins during two different sessions, averaged over all pairs of sessions and all animals.

To quantify the consistency of the population dynamics across trials (figure 5a-c), population vectors were computed by averaging neural activity over time or location bins within each trial, as described above. Then a similar correlation matrix was constructed where each element is the cosine similarity between a pair of population vectors from different trials.

To test that the matrix shows a significant diagonal pattern, neural activity across all bins within each session (Figure 5d-f) or trial (Figure 5a-c) was shuffled 10000 times independently for each neuron and matrices from this shuffled data were constructed. We characterized the degree to which each matrix shows a diagonal pattern by an index *d*, which equals the difference between the average value of the near-diagonal matrix elements to that of the off-diagonal matrix elements. The near-diagonal matrix elements are those whose row and column indices are differed by less than half the dimension of the matrix.

Mathematically, 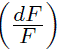, where *N* is the dimension of the matrix and indicates the mean. Then we counted how many matrices constructed from the shuffled data have a value *d* greater than the matrix obtained from the true data. As a result, none of the 10000 matrices from the shuffled data has a higher *d* than the matrices in Figure 5.

The same method was used to quantify the significance of the diagonal patterns of the matrices obtained from the decoding analyses (Figure 6), except that the shuffling was done 1000 times.

### Firing consistency score

To assess the consistency of the single neuron dynamics across repeated trials or sessions, we computed a firing consistency score for each neuron. For each cell *n* and each pair of sessions or trials (for example *i* and *j*), we computed the Pearson correlation coefficient between the coarse-grained activity vectors *v_n,i_* and *v_n,j_* obtained from the method described above. Then we shuffled the entries in each activity vector and computed the Pearson’s correlation coefficient again. This was repeated for 100 times. The Pearson correlation coefficients from all pairs of sessions (or trials) were then averaged to obtain a mean Pearson correlation coefficient across pairs of sessions, for the true data and each shuffle. For the second-level scores (Figure 4a-c), the Pearson correlation coefficients were only computed between pairs of trials with the same trial type. The firing consistency score was computed as the percentile where the true mean Pearson correlation is at among all the shuffles (if there are multiple shuffles that yield the same Pearson’s correlation as the true data, the median percentile was used). Sessions or trials where the neuron does not have any calcium transient event during the selected time period were excluded from the analysis.

### Firing linearity score

To disentangle the gradually ramping/decaying activity from more complex temporal modulations, we computed a firing linearity score for each neuron. For a given neuron *n* and session (or trial) *i*, we fitted a linear model as a function of the bin number for the coarse-grained activity *v_n,i_* of that neuron. The F-statistic of this linear model was computed along with those obtained from 100 shuffled activity vectors (shuffling was performed in the same way as in computing the firing consistency score). The F-statistics for both the true and shuffled data were then averaged across all sessions (or trials) to obtain a mean F-statistic for the true data and for each shuffle. The firing linearity score was computed as the percentile of the true mean F-statistic among all the shuffled mean F-statistic was computed (if the F statistic of multiple shuffles equal the true F statistic, the median percentile was used). For each neuron, sessions (or trials) where no calcium transients were observed were excluded from the analysis.

### Temporal information score

To assess the single cell basis for the coding of trial number, the temporal information metric, which was used as a criteria for identifying hippocampal “time cells” (Mau et al., 2018), was adapted to identify cells that robustly carry information about trial number during a session. First, the average activity during each trial bin was computed (see “Coarse-graining for the across session dynamics”. The zscore step was not performed), and further averaged across sessions. This results in a time series of length *N* where *N* is the number of trial bins (*N* = 5 in our analysis). *v_n_* = *v_n,_*_1_*, v_n,_*_2_*, v_n,_*_3_*, v_n,_*_4_*, v_n,_*_5_ . Then the temporal information for neuron *n* is the negative Shannon entropy of the normalized time series

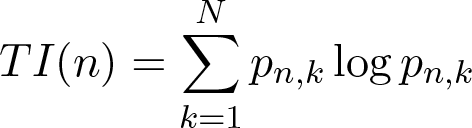

where 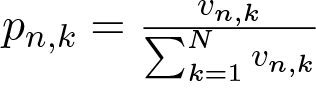.

If a cell is n*^k^*o^=^t^1^ modulated by trial number, or it is modulated in a way that is not consistent across sessions, its session-averaged activity *v_n,_*_1_*, v_n,_*_2_*, v_n,_*_3_*, v_n,_*_4_*, v_n,_*_5_ would be weakly modulated by trial number. Since the Shannon entropy is the largest for uniform probability distributions, the TI (which is the negative Shannon entropy) would be small. The TI was then compared with the TIs of 1000 shuffles where neural activity was randomly shuffled across trial bin number before averaging over sessions, and the percentile of the true TI among all the shuffled was defined as the “temporal information score”.

## Data availability

The data that support the findings of this study are available from the corresponding author upon reasonable request.

## Supplementary figures

**Figure S1.**
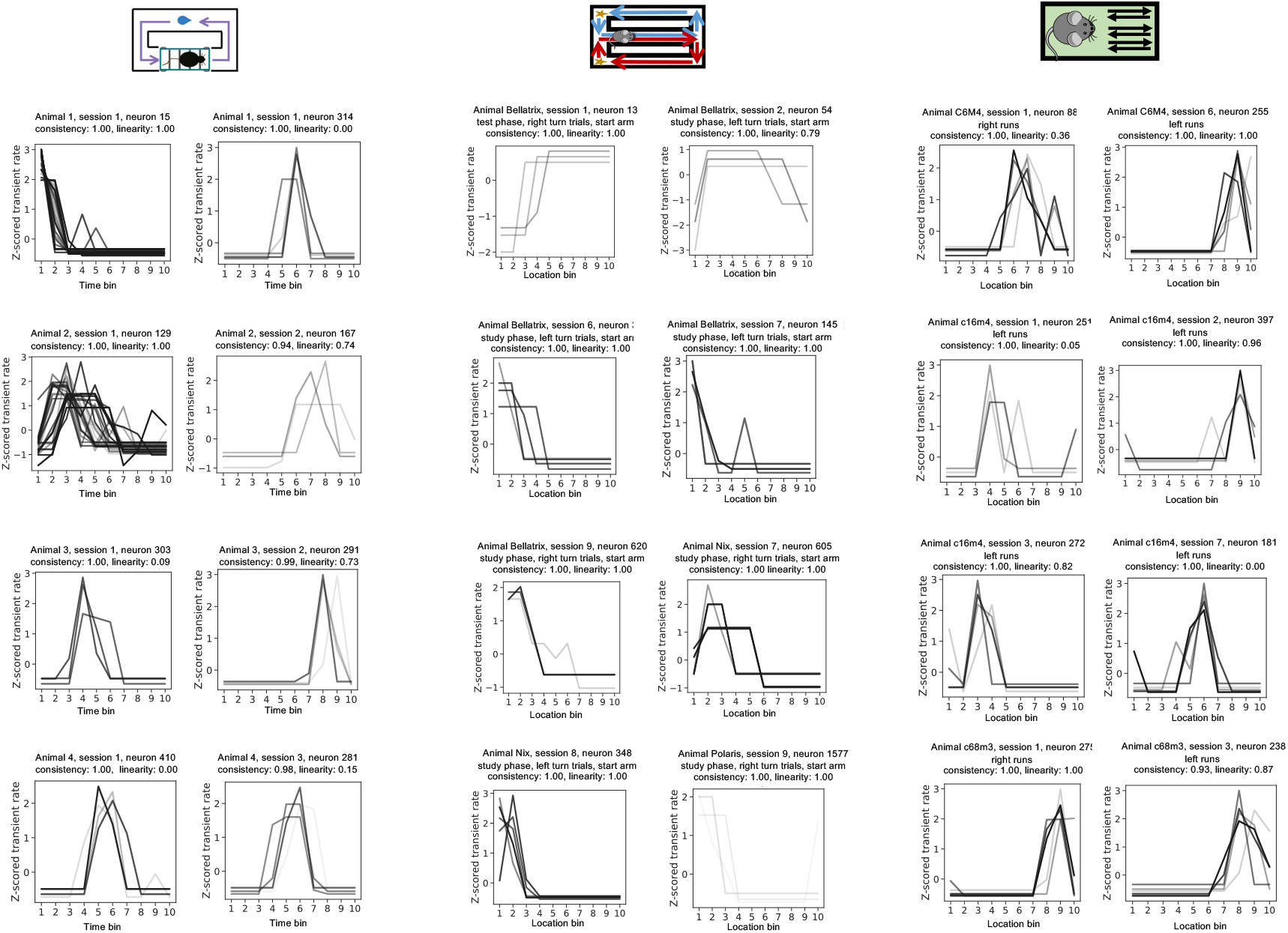
Additional hippocampal neurons that fire consistently across trials. Each line represents the z-scored transient rate of that neuron during a trial. Darker lines indicate earlier trials within a session. Trials where the neuron is not identified or is inactive are not plotted.

**Figure S2.**
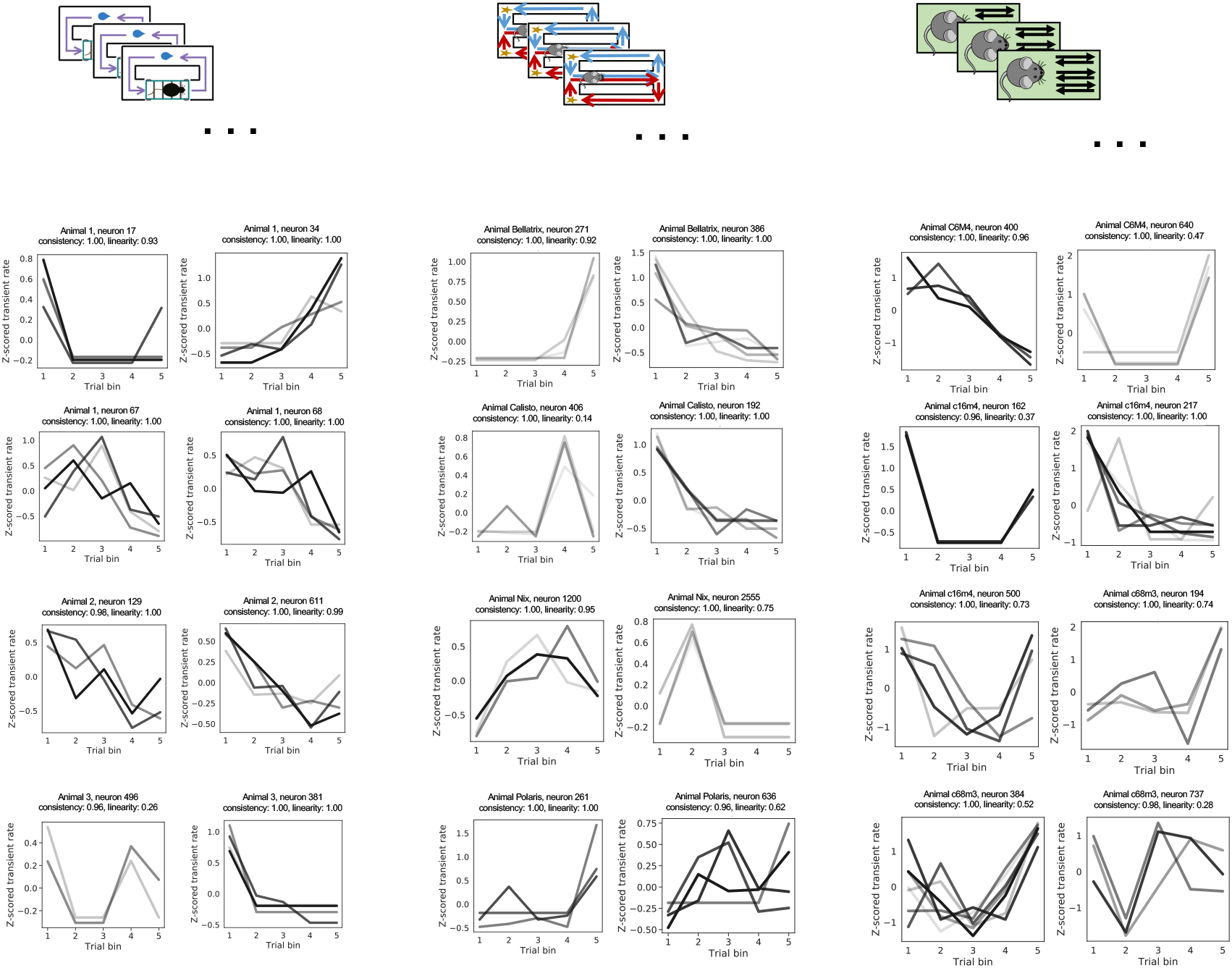
Additional hippocampal neurons that fire consistently across multiple sessions. Each line represents the z-scored transient rate of that neuron during a session. Darker lines indicate earlier sessions. Sessions where the neuron is not identified or is inactive are not plotted.

**Figure S3.**
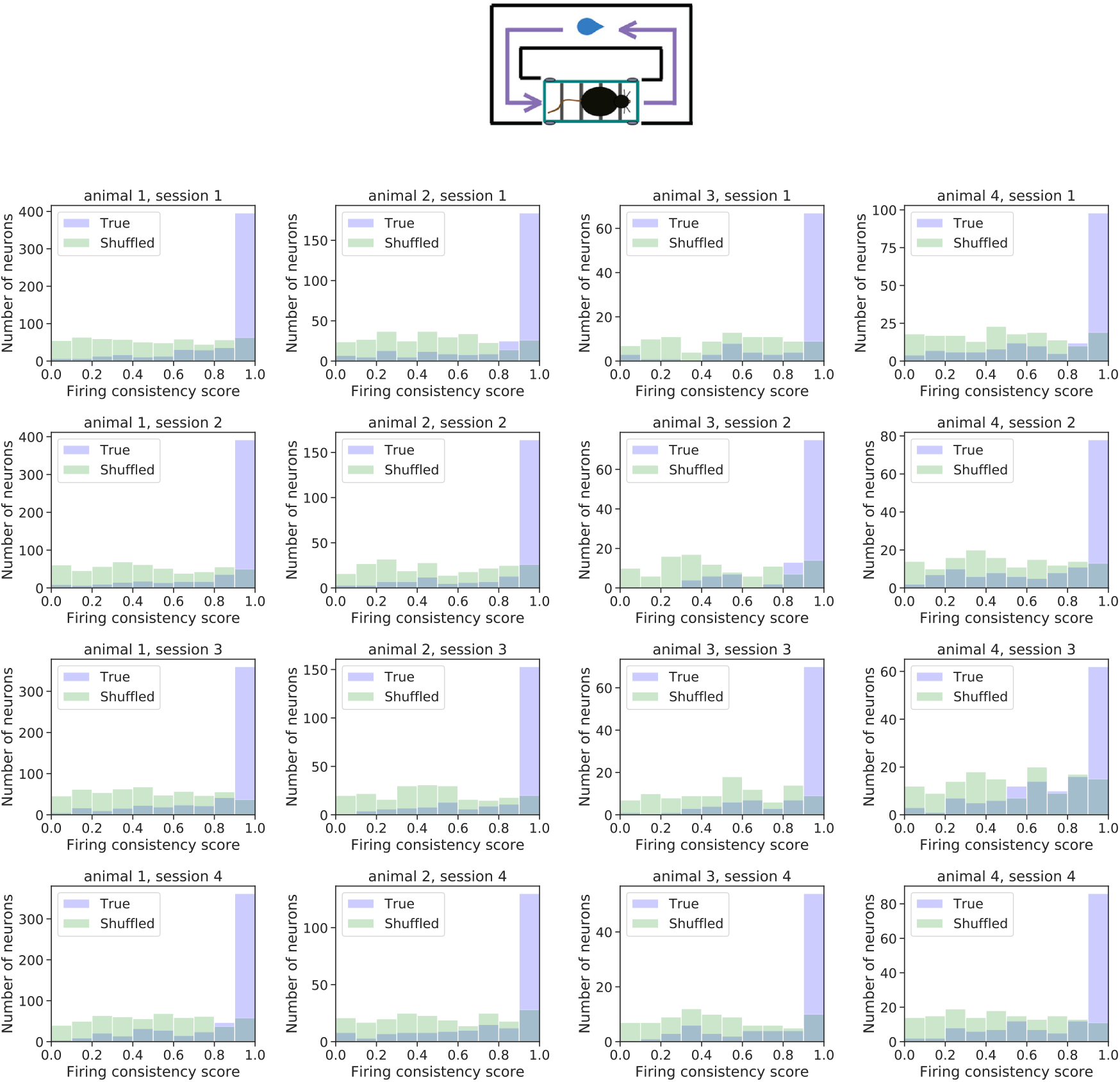
The distribution of the across-trial firing consistency score for each individual session in the treadmill running task.

**Figure S4.**
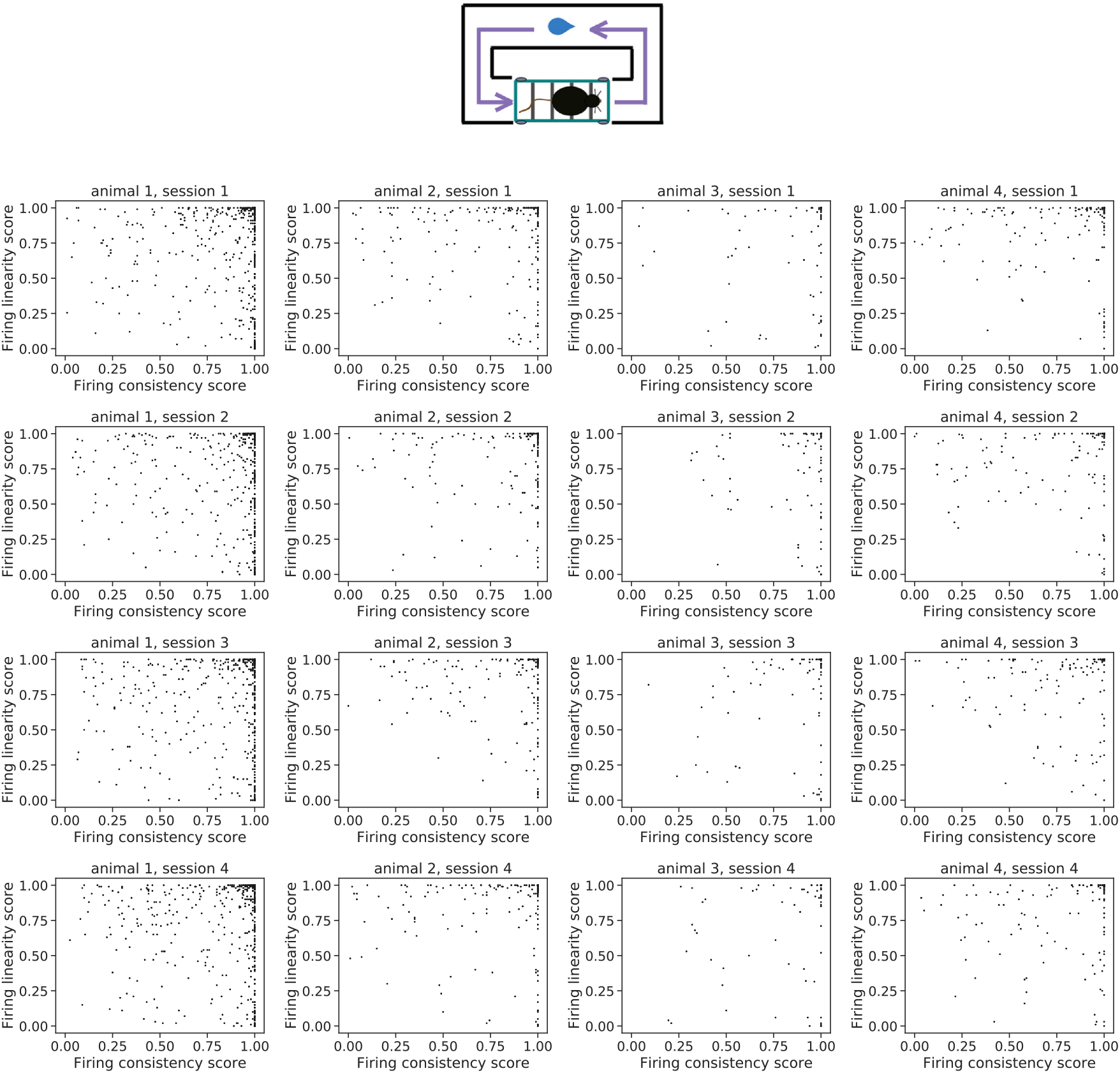
The across-trial firing consistency score plotted against the across-trial firing linearity score for each individual session in the treadmill running task.

**Figure S5.**
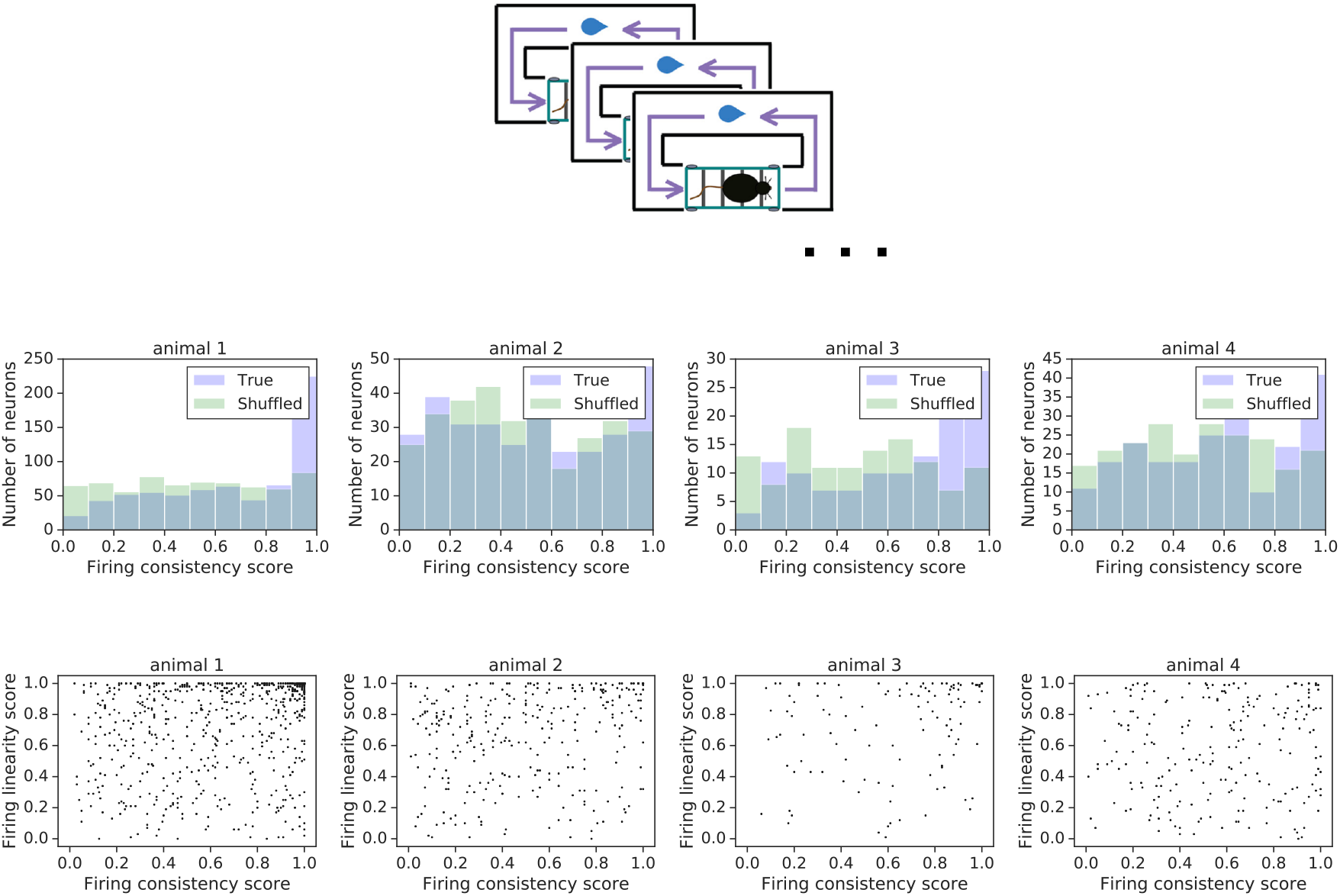
The distribution of the across-session firing consistent score for real and shuffled data (top) and he joint distribution of the across-session firing consistency score and firing linearity score (bottom) for each individual mouse in the treadmill running task.

**Figure S6.**
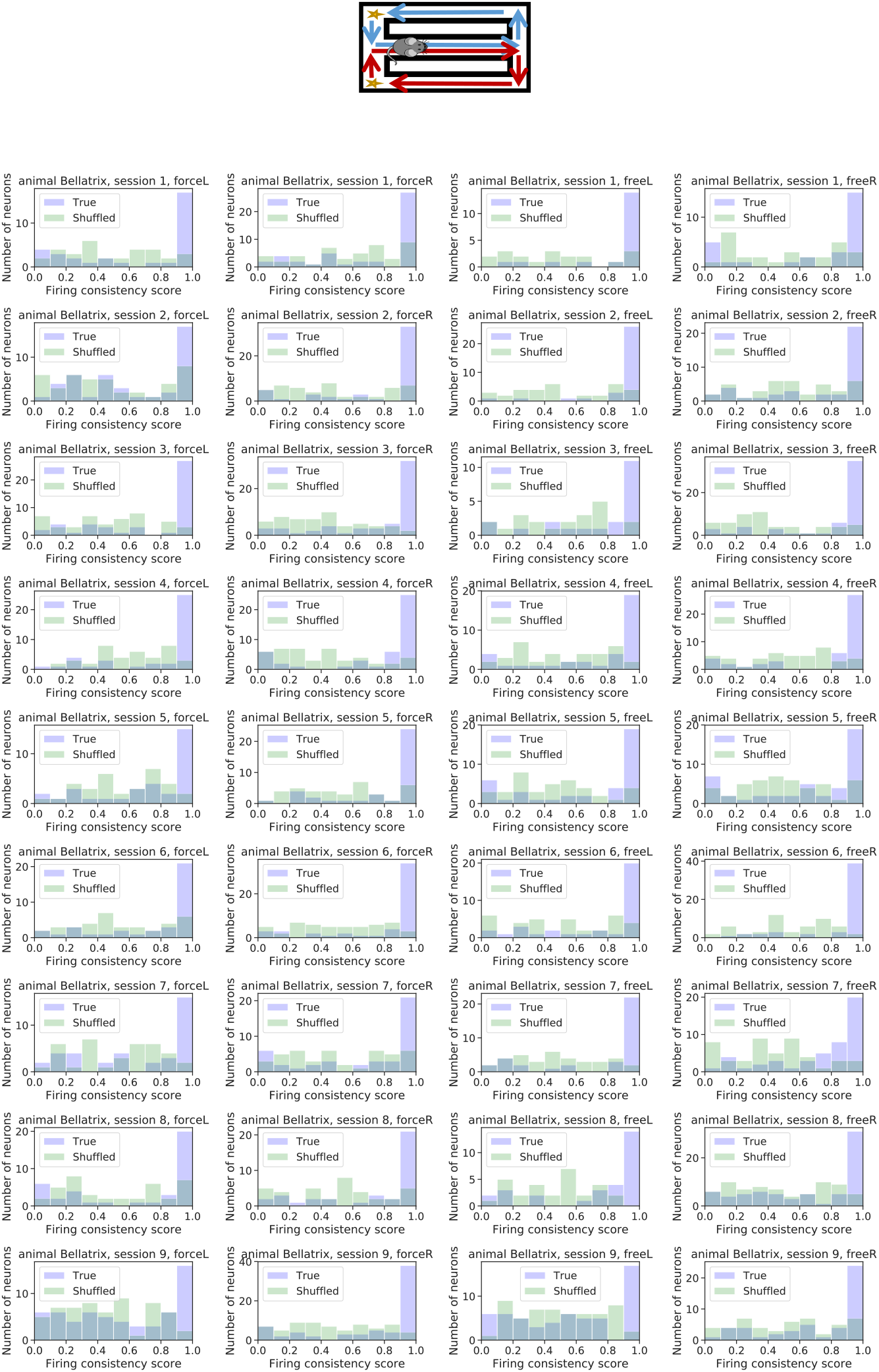
The distribution of the across-trial firing consistent score for real and shuffled data for each individual session and trial type in the spatial alternation task. Data for mouse Bellatrix.

**Figure S7.**
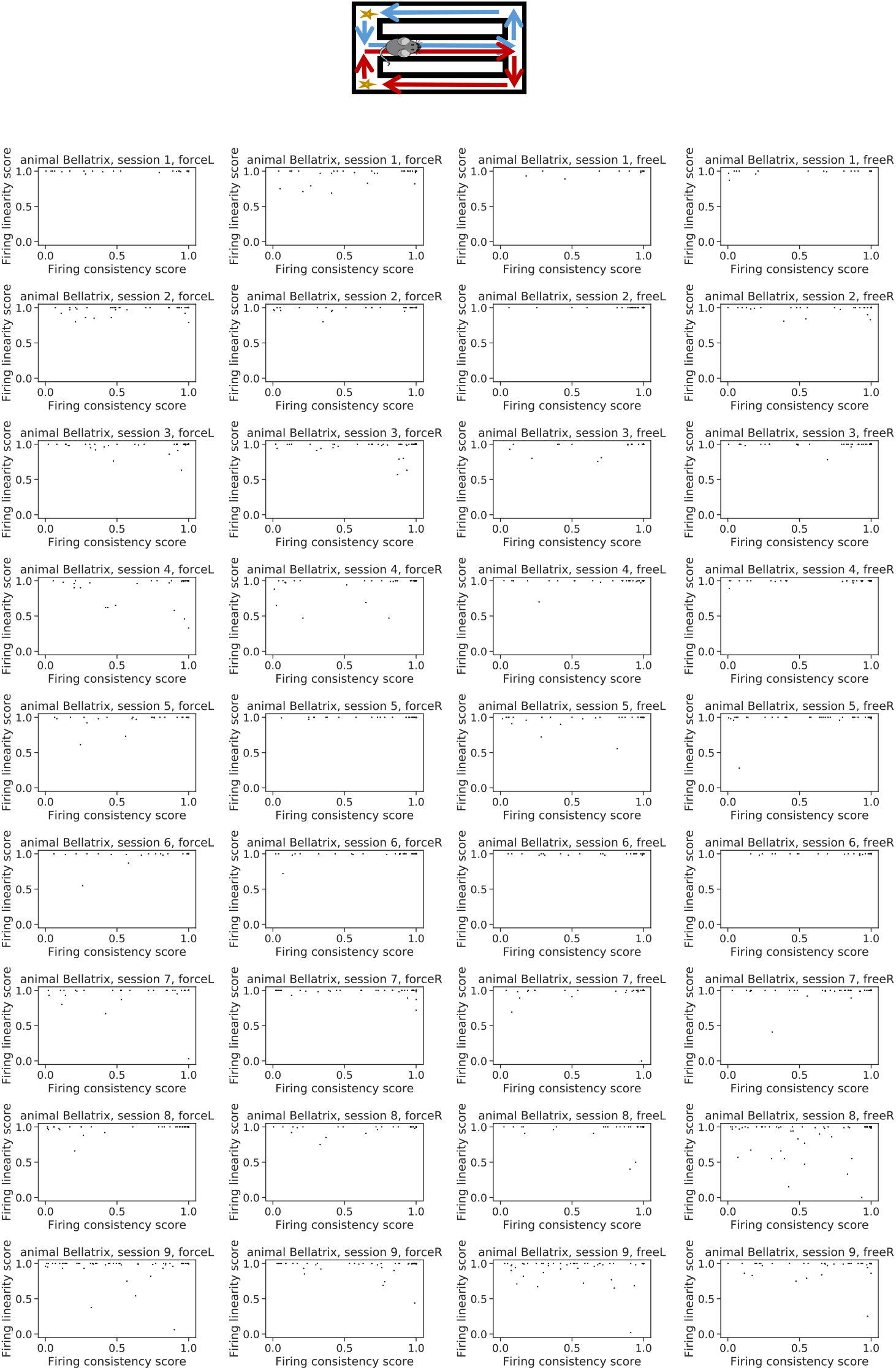
The joint distribution of the across-trial firing consistency score and firing linearity score for each individual session and trial type in the spatial alternation task. Data for mouse Bellatrix.

**Figure S8.**
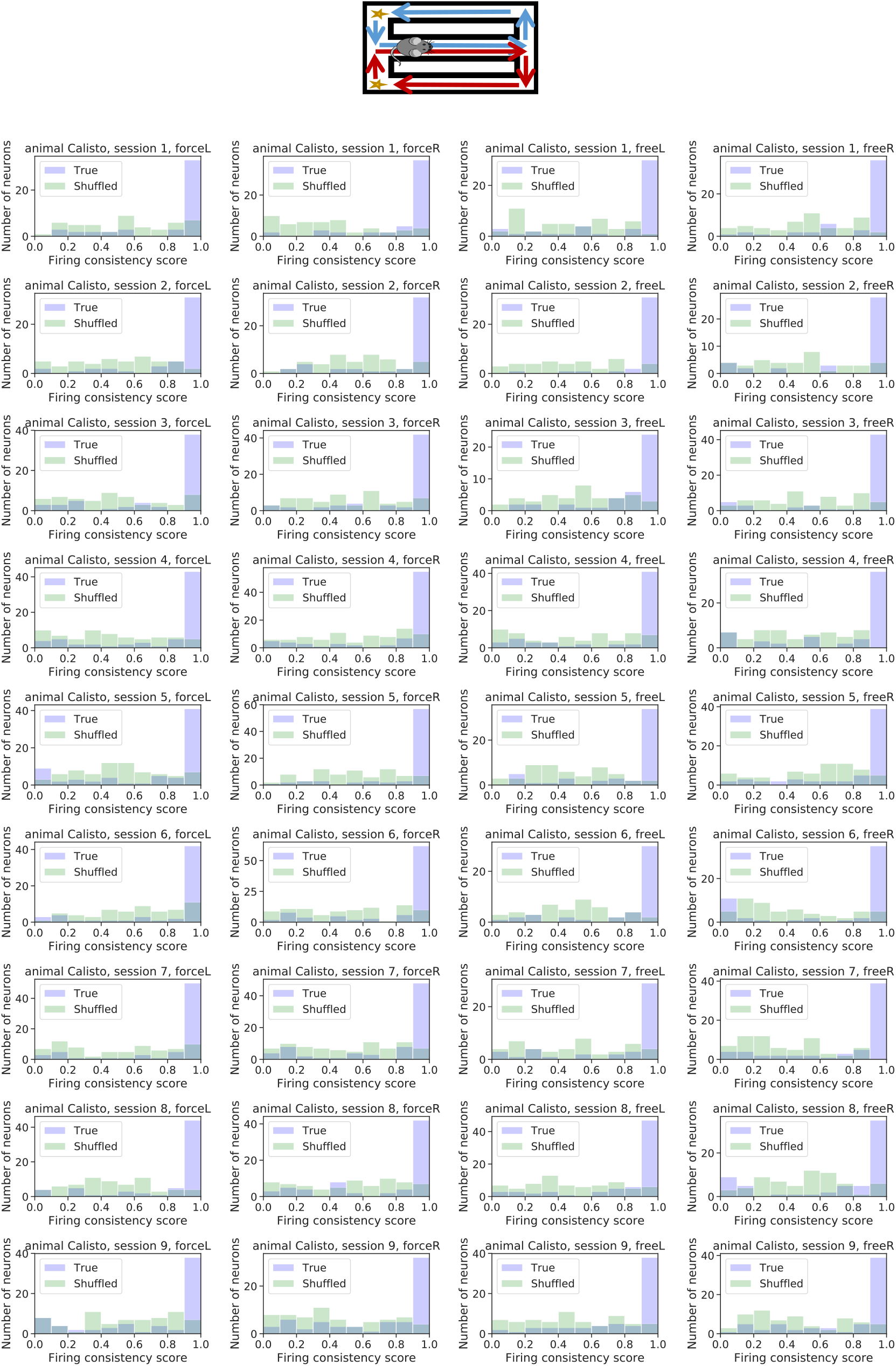
The distribution of the across-trial firing consistent score for real and shuffled data (left) for each individual session and trial type in the spatial alternation task. Data for mouse Calisto.

**Figure S9.**
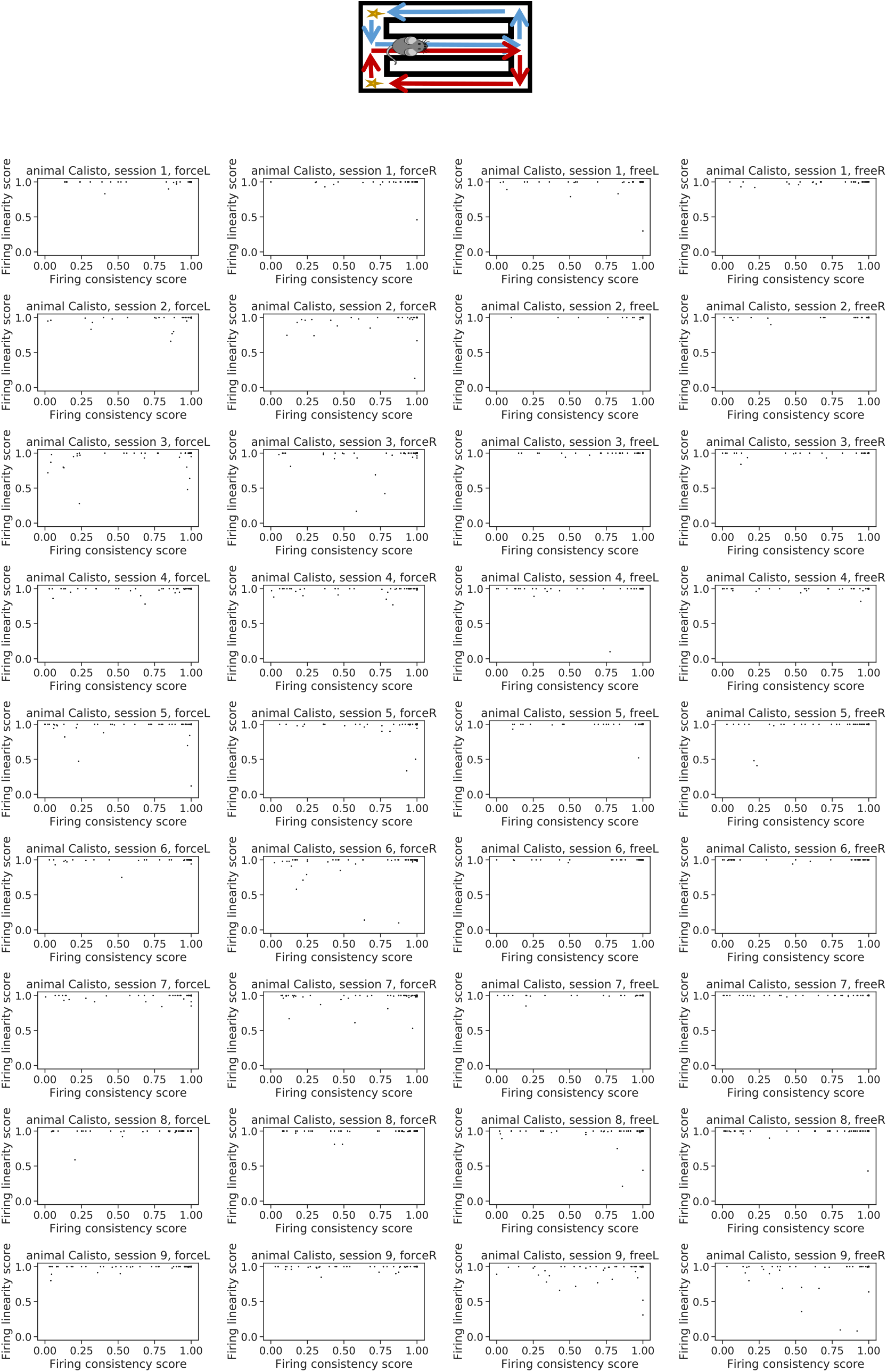
The joint distribution of the across-trial firing consistency score and firing linearity score for each individual session and trial type in the spatial alternation task. Data for mouse Calisto.

**Figure S10.**
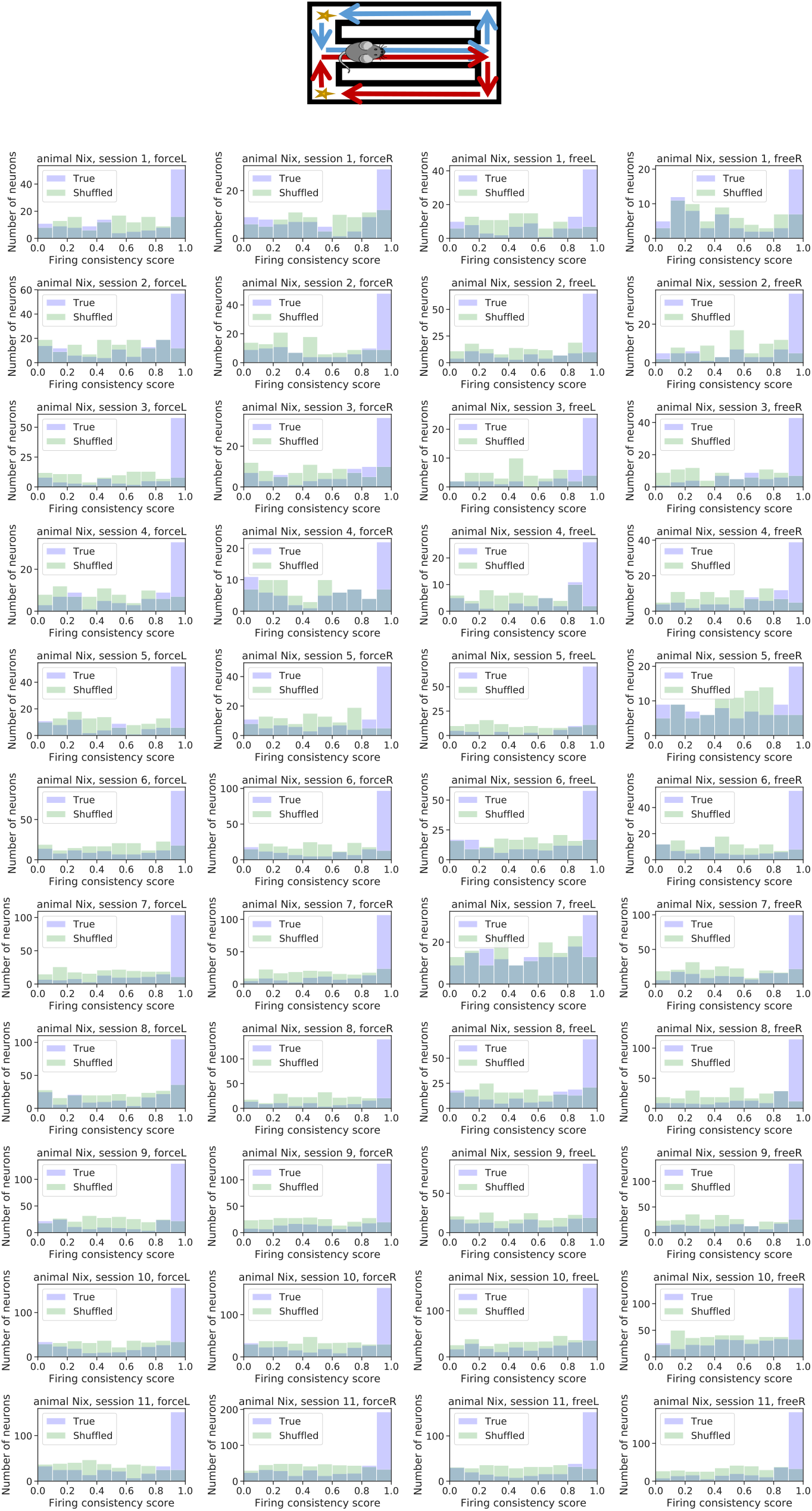
The distribution of the across-trial firing consistent score for real and shuffled data. Data for mouse Nix.

**Figure S11.**
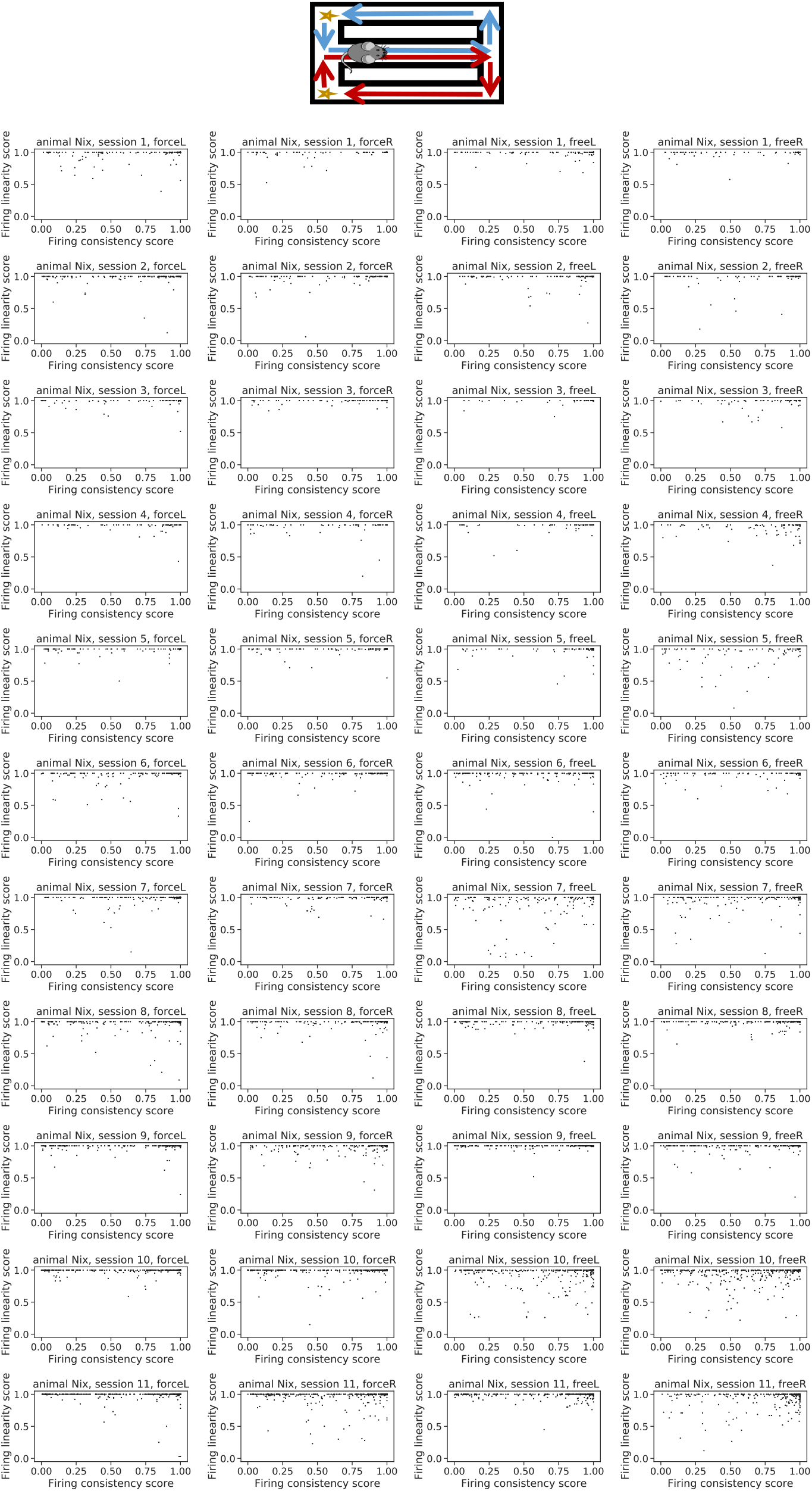
The joint distribution of the across-trial firing consistency score and firing linearity score for each individual session and trial type in the spatial alternation task. Data for mouse Nix.

**Figure S12.**
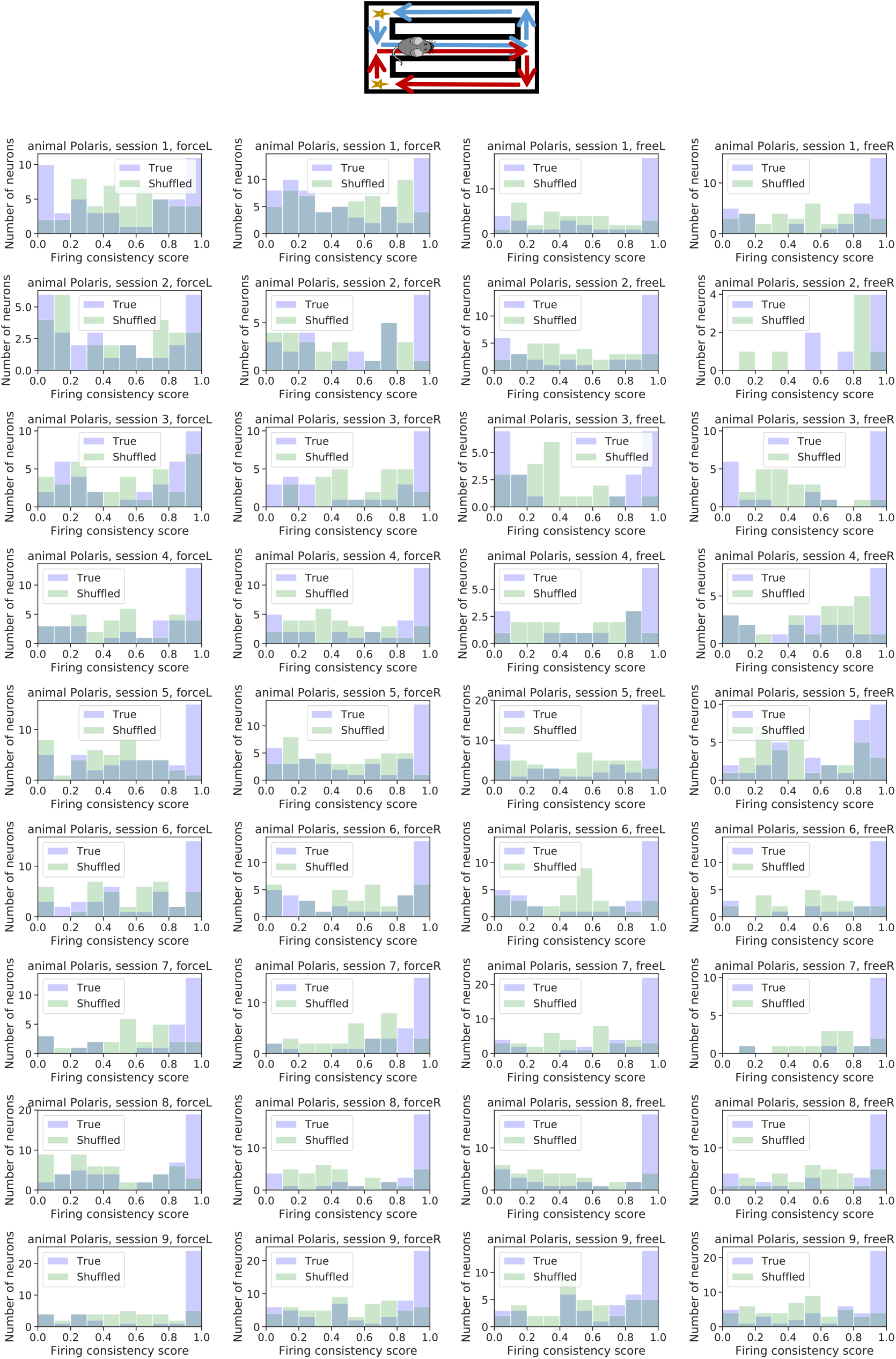
The distribution of the across-trial firing consistent score for real and shuffled data. Data for mouse Polaris.

**Figure S13.**
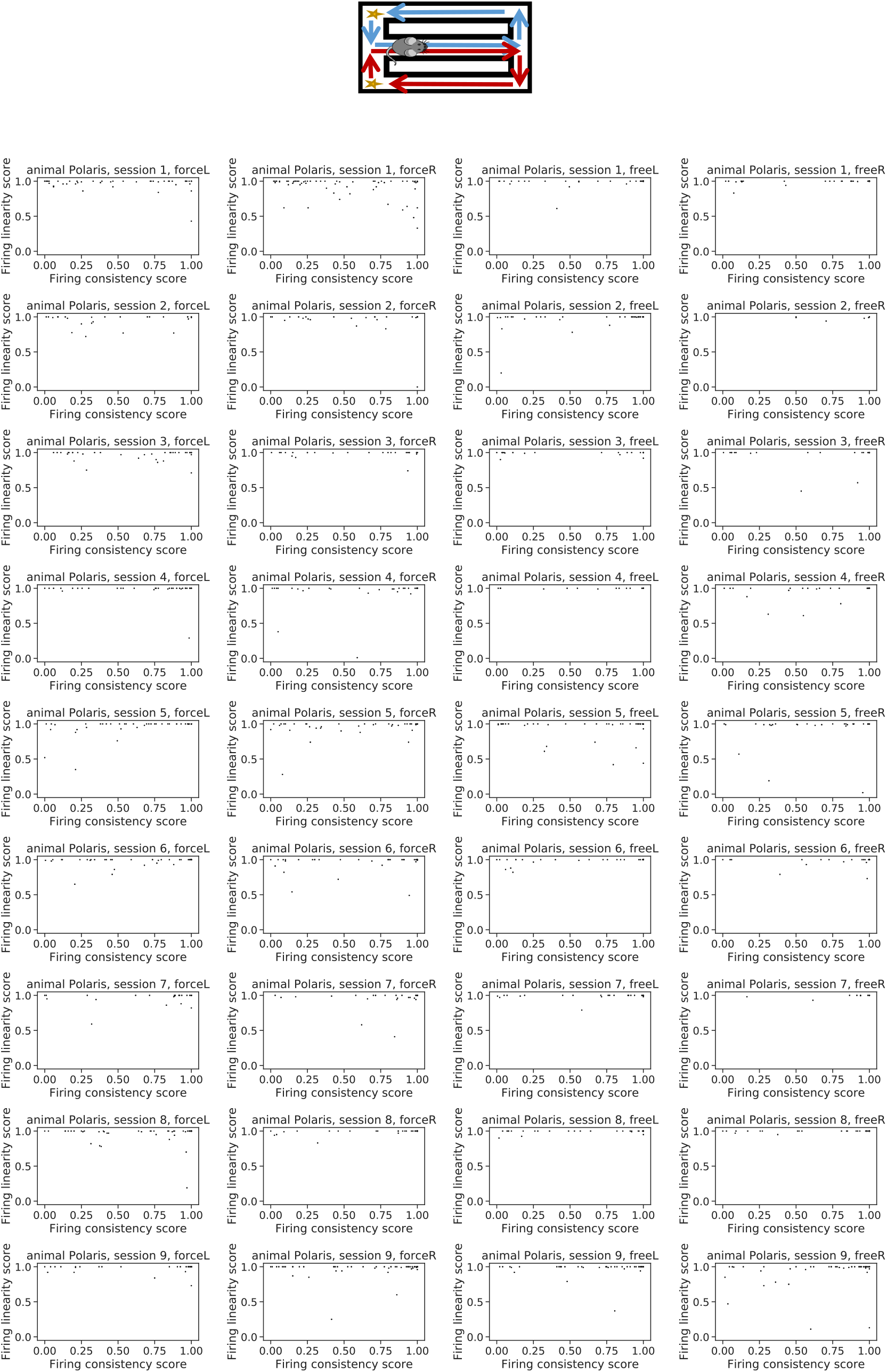
The joint distribution of the across-trial firing consistency score and firing linearity score for each individual session and trial type in the spatial alternation task. Data for mouse Polaris.

**Figure S14.**
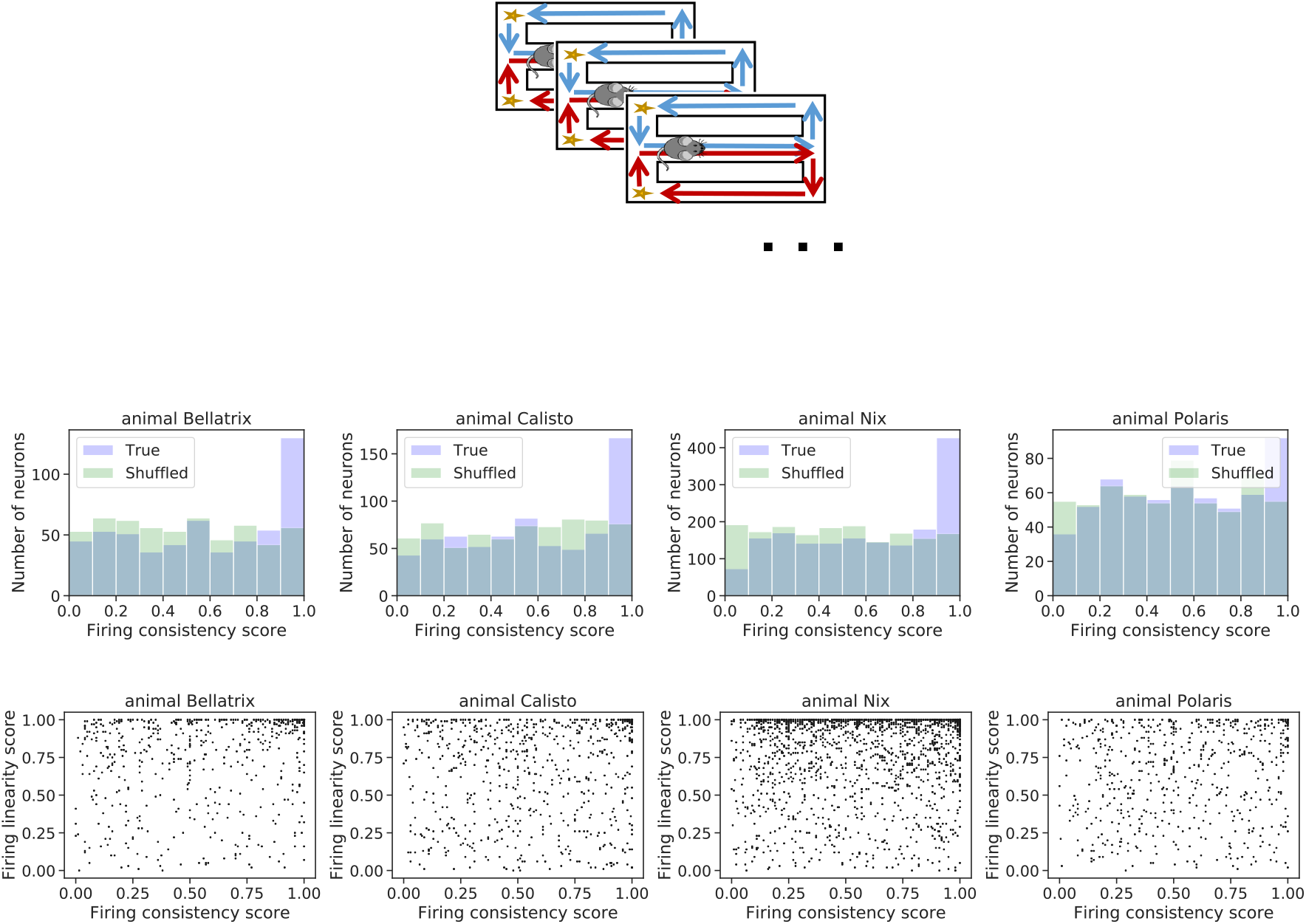
The distribution of the across-session firing consistent score for real and shuffled data (top) and the joint distribution of the across-session firing consistency score and firing linearity score (bottom) for each individual mouse in the spatial alternation task.

**Figure S15.**
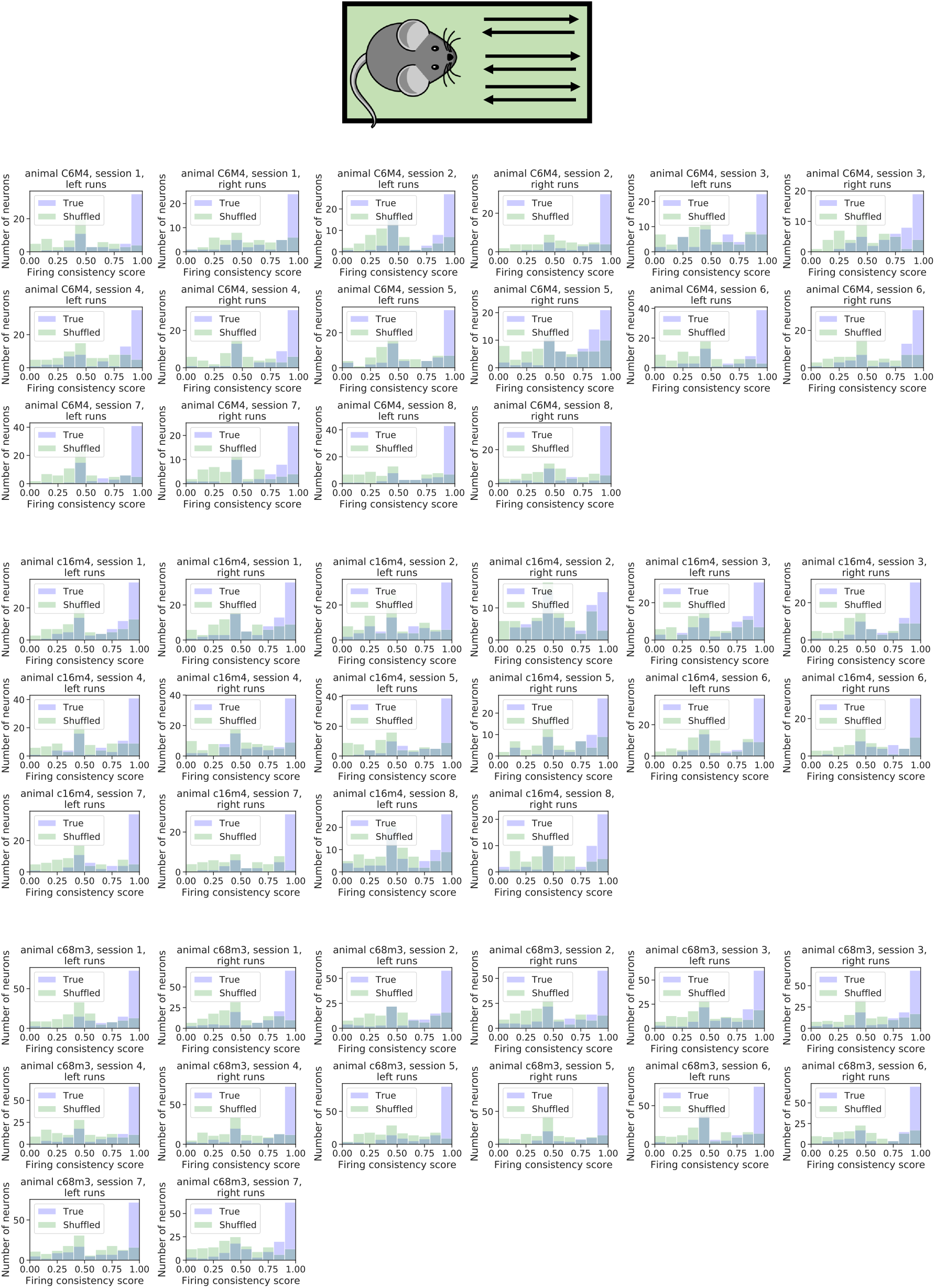
The distribution of the across-trial firing consistency score for each individual session in the linear track task. Data shown separately for left and right runs.

**Figure S16.**
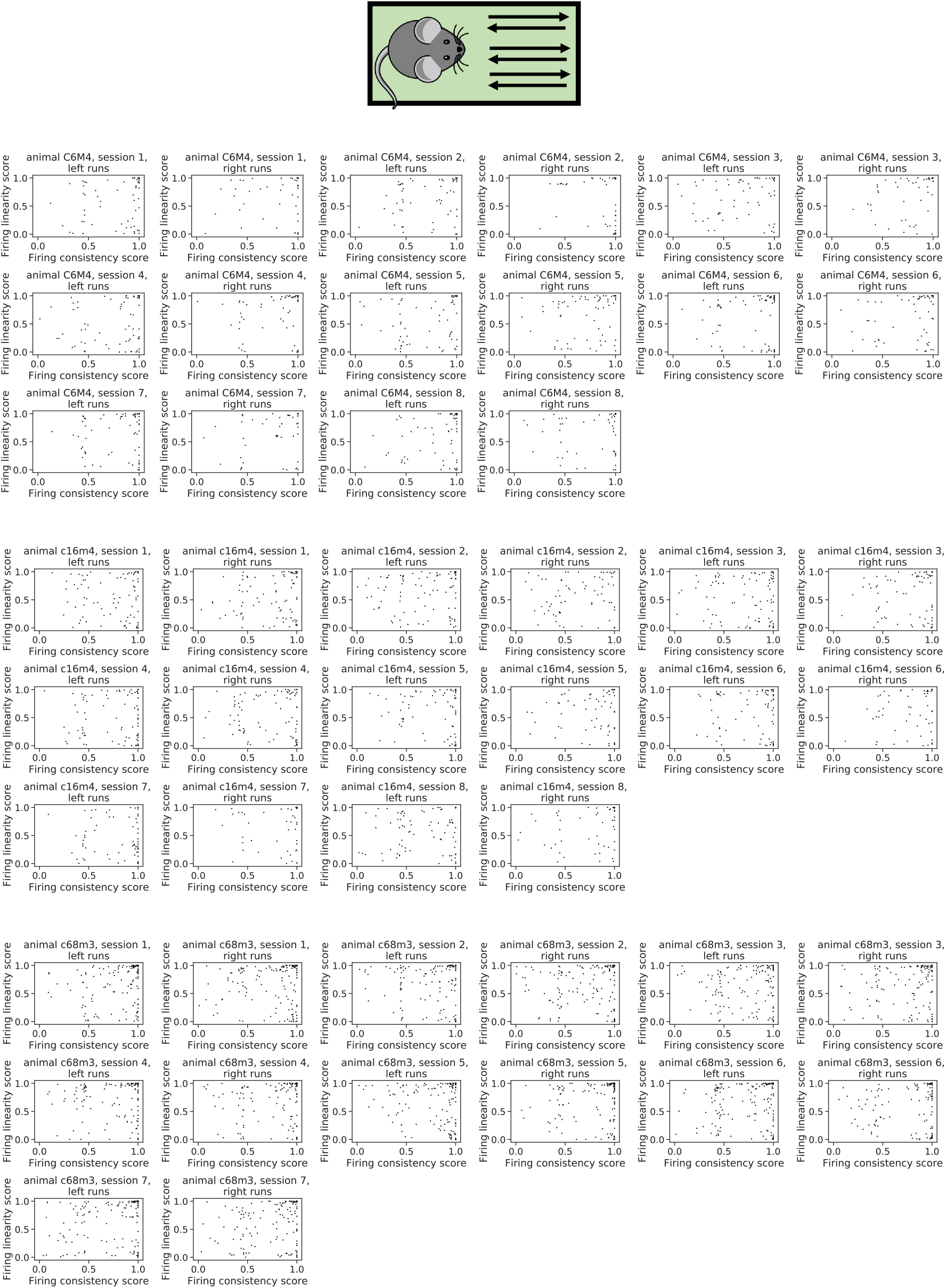
The across-trial firing consistency score plotted against the across-trial firing linearity score for each individual session in the linear track task.

**Figure S17.**
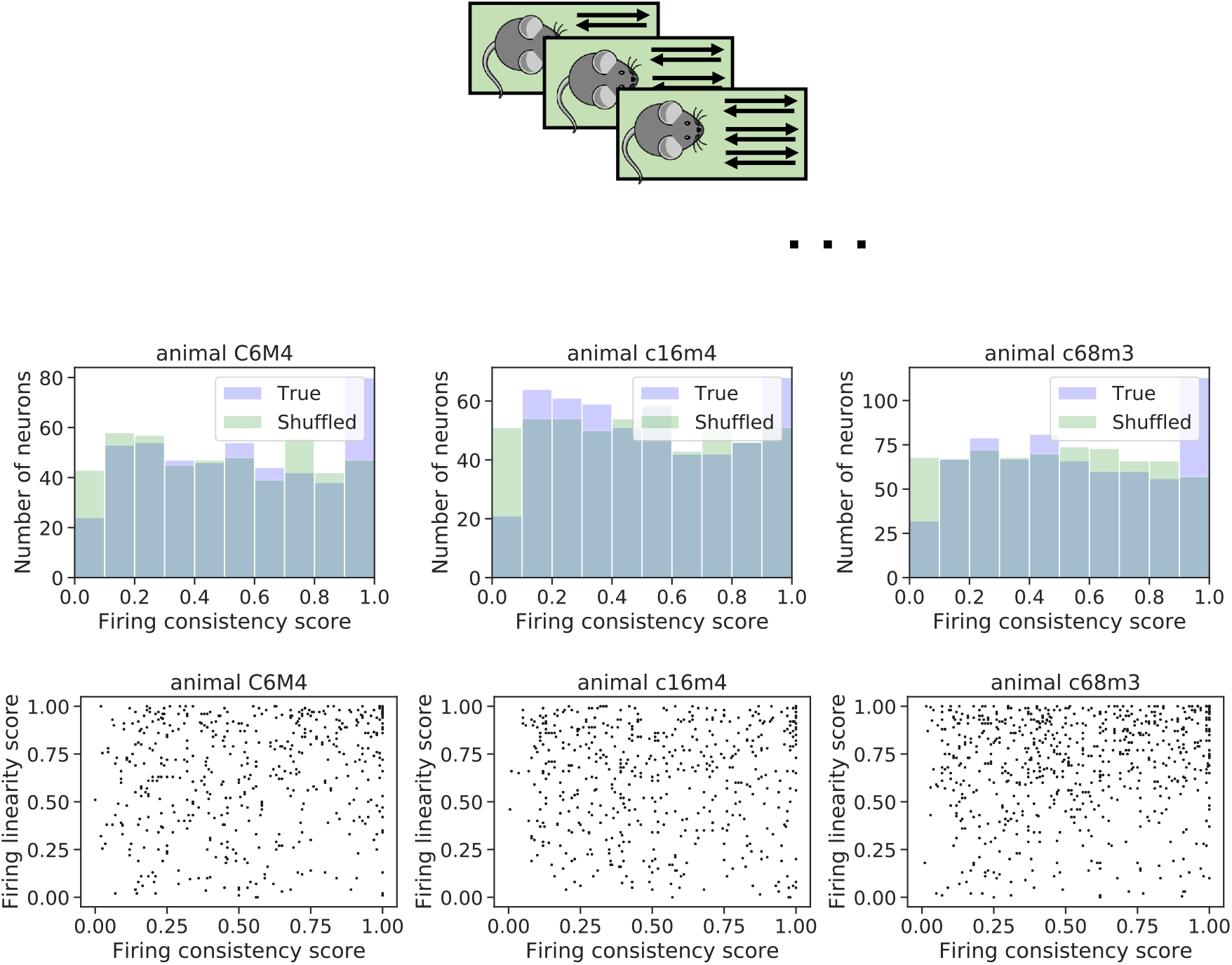
The distribution of the across-session firing consistent score for real and shuffled data (top) and the joint distribution of the across-session firing consistency score and the firing linearity score (bottom) for each individual mouse in the linear track task.

**Figure S18.**
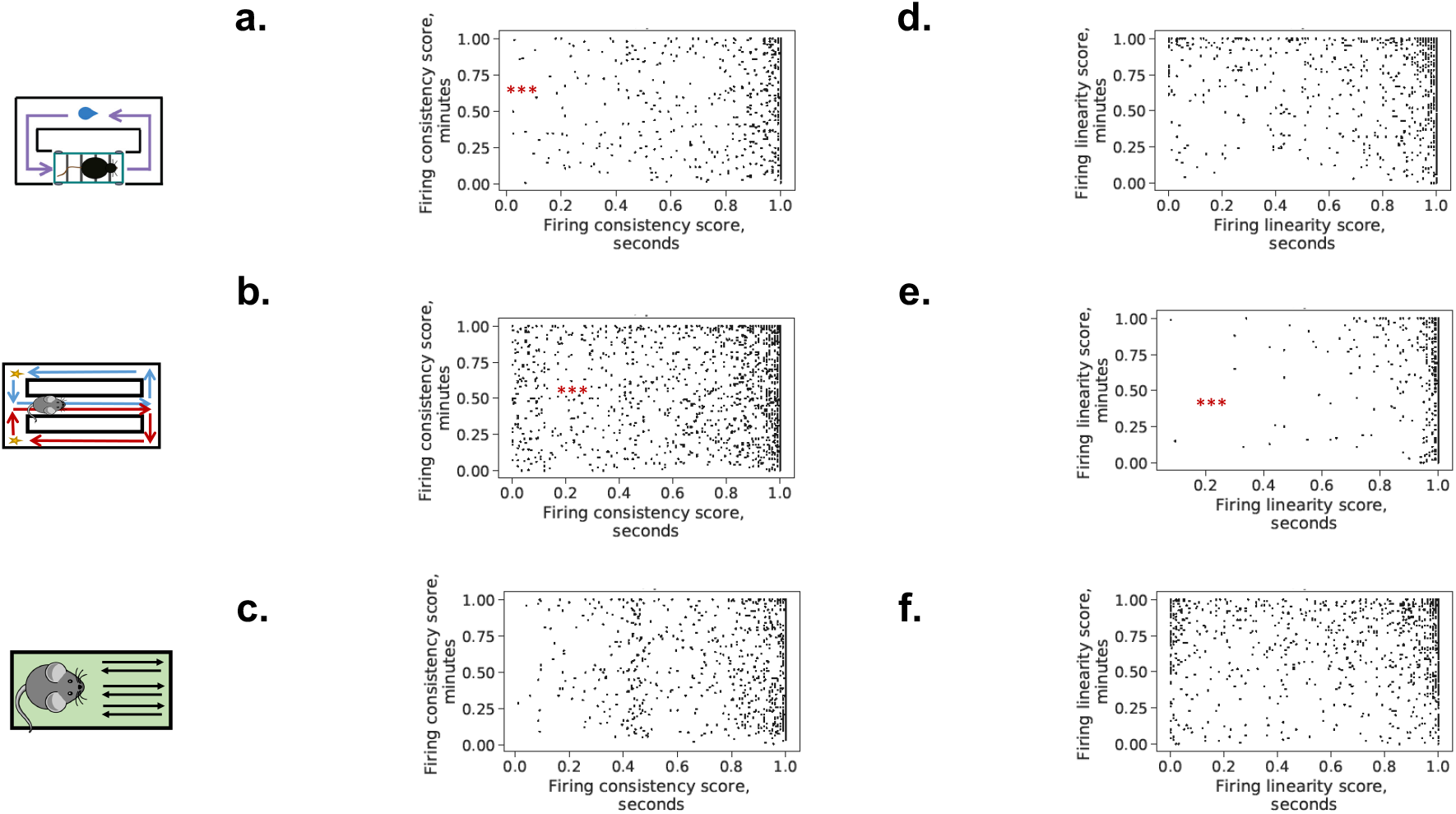
The correlation between the firing consistency score on the timescales of seconds and minutes for the treadmill running (a), spatial alternation (b) and linear track (c) experiments. The same for the firing linearity score (d-f). Kendall’s *τ*: **a**: *τ* (1245) = 0.13*, p <* 10*^−^*^8^. **b**: *τ* (3019) = 0.079*, p <* 10*^−^*^8^. **c**: *τ* (1099) = −0.01*, p* = 0.6. **d**: *τ* (1245) = 0.014*, p* = 0.5. **e**: *τ* (3019) = 0.12*, p <* 10*^−^*^15^. **f**: *τ* (1099) = 0.011*, p* = 0.6.

**Figure S19.**
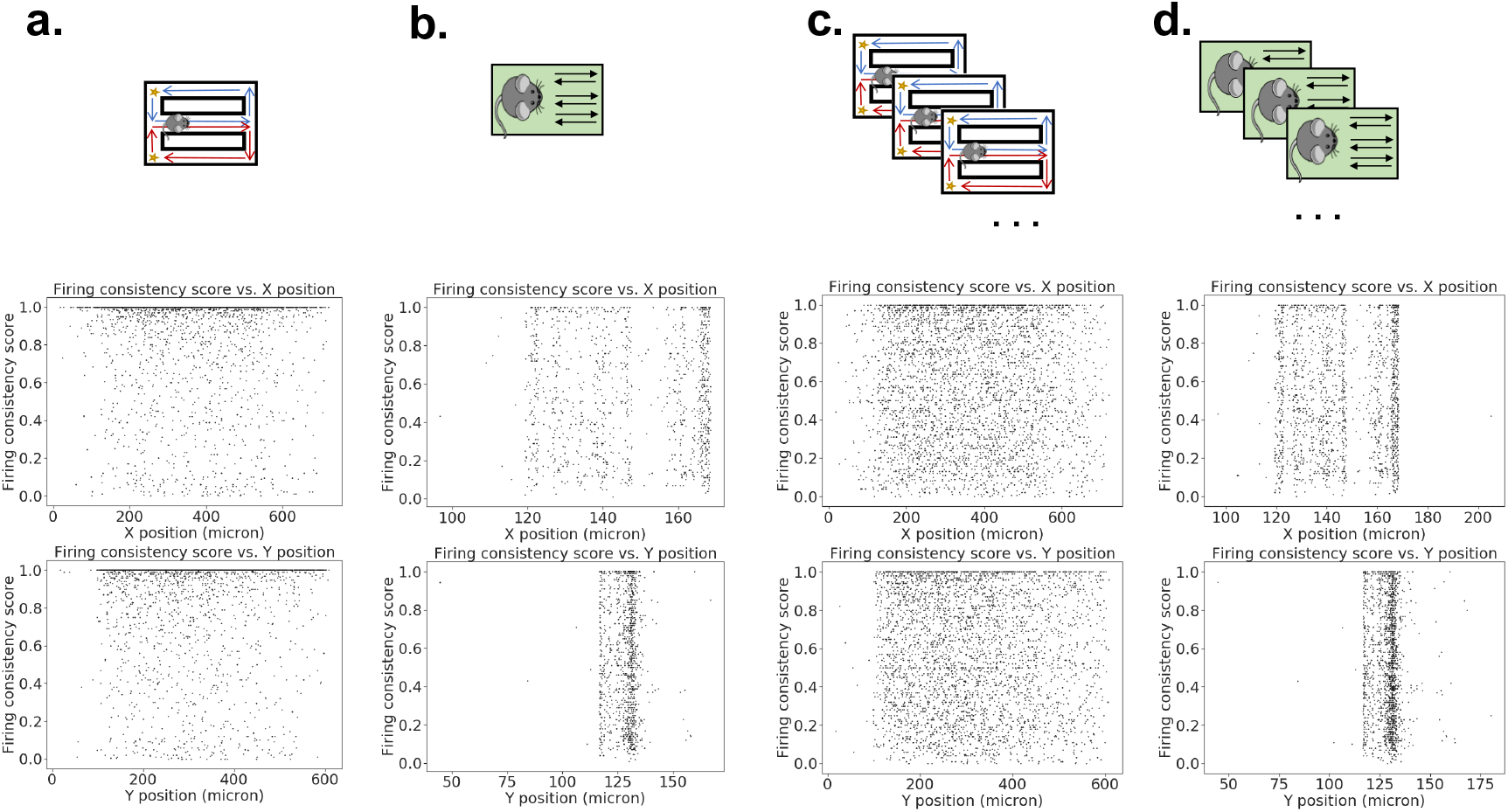
Firing consistency ranks are not significantly correlated with x or y position in the field of view, either on the timescale of seconds (a, b), or on the timescale of minutes (c, d). A statistical test using Pearson’s correlation coefficient was conducted. **a** top: *r* = − 0.02, *p* = 0.12; bottom: *r* = 0.0004, *p* = 0.98. **b** top: *r* = −0.01*, p* = 0.68; bottom: *r* = −0.02*, p* = 0.43. **c** top: *r* = 0.003, *p* = 0.84; bottom: *r* = −0.02, *p* = 0.23. **d** top: *r* = 0.01*, p* = 0.67; bottom: *r* = −0.03*, p* = 0.14.

**Figure S20.**
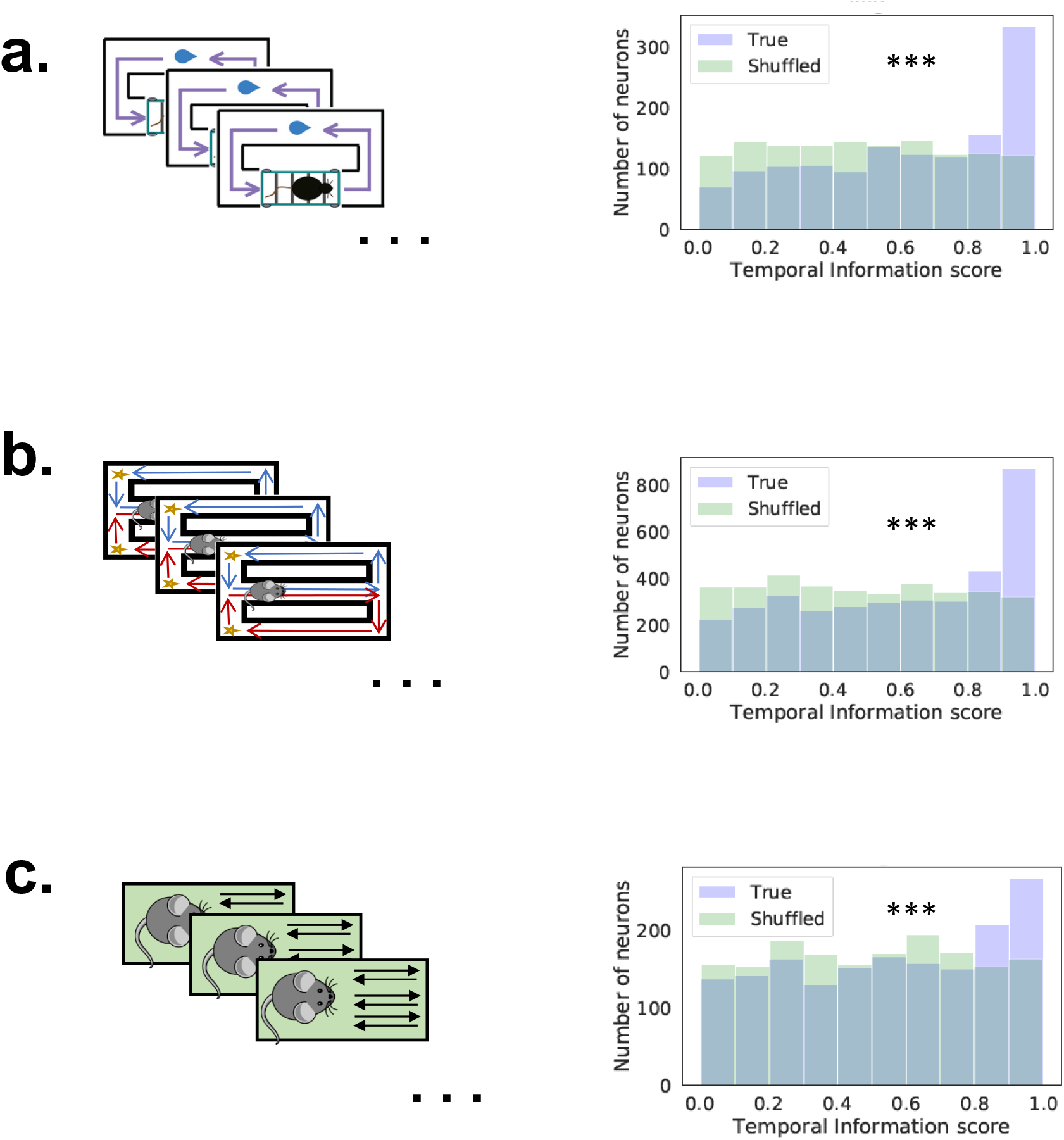
The distributions of the temporal information score are significantly biased towards larger values compared with shuffled data. See Methods section for the definition of the temporal information score.

**Figure S21.**
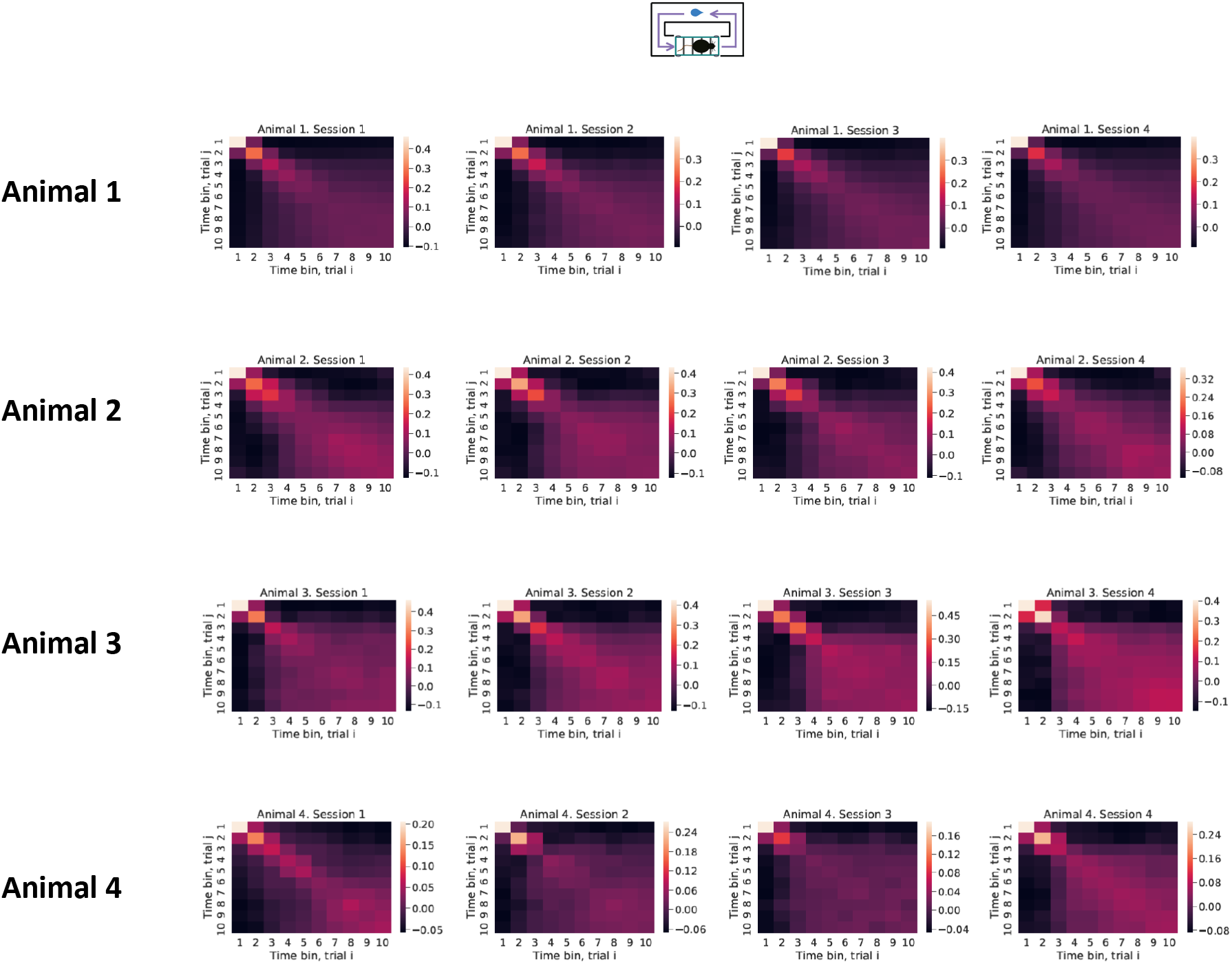
The cross-trial correlations for each individual session in the treadmill running task.

**Figure S22.**
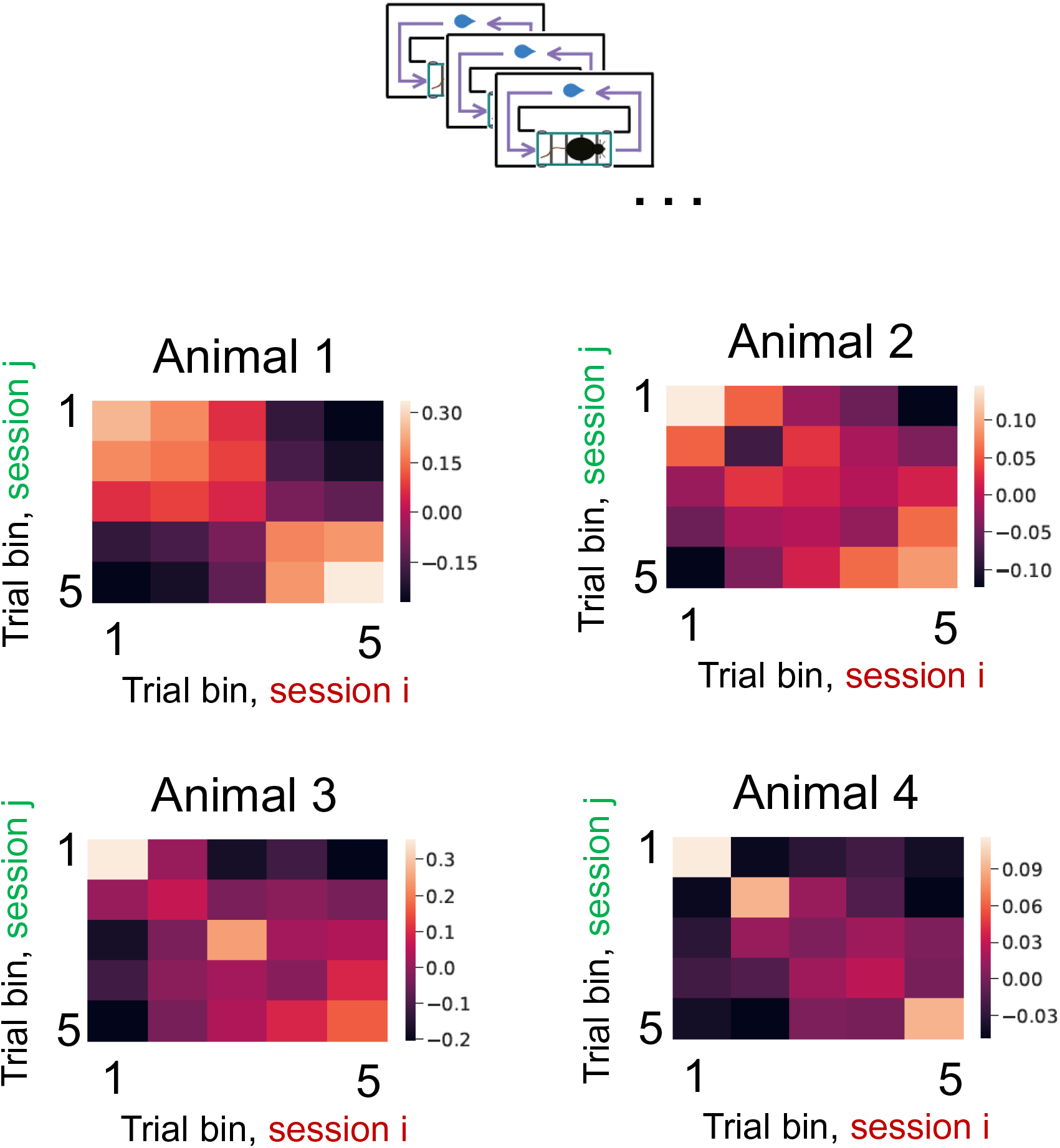
The cross-session correlations for each individual mouse in the treadmill running task.

**Figure S23.**
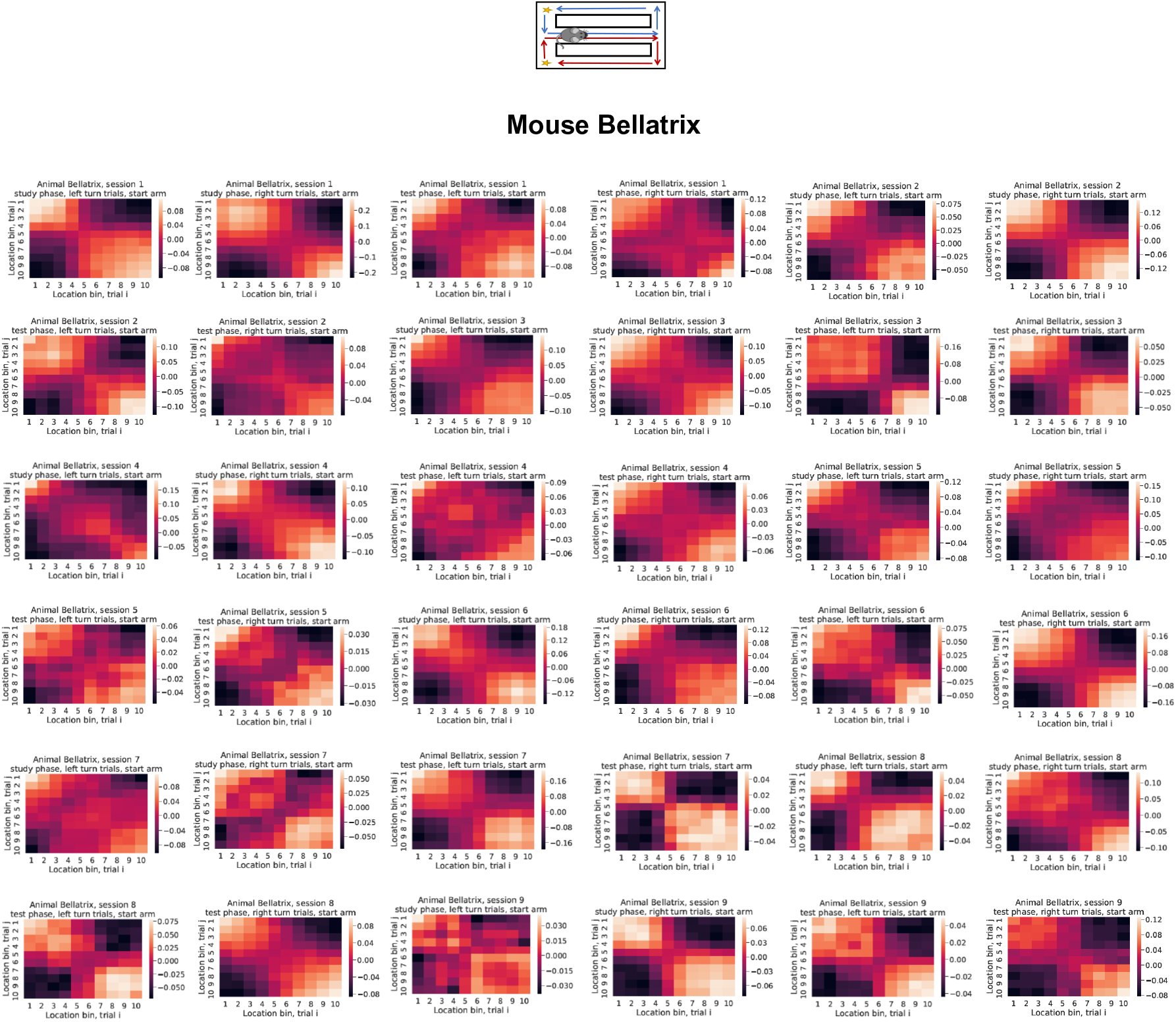
The cross-trial correlations for each individual session, task phase and turn direction in the spatial alternation task. Data for mouse Bellatrix.

**Figure S24.**
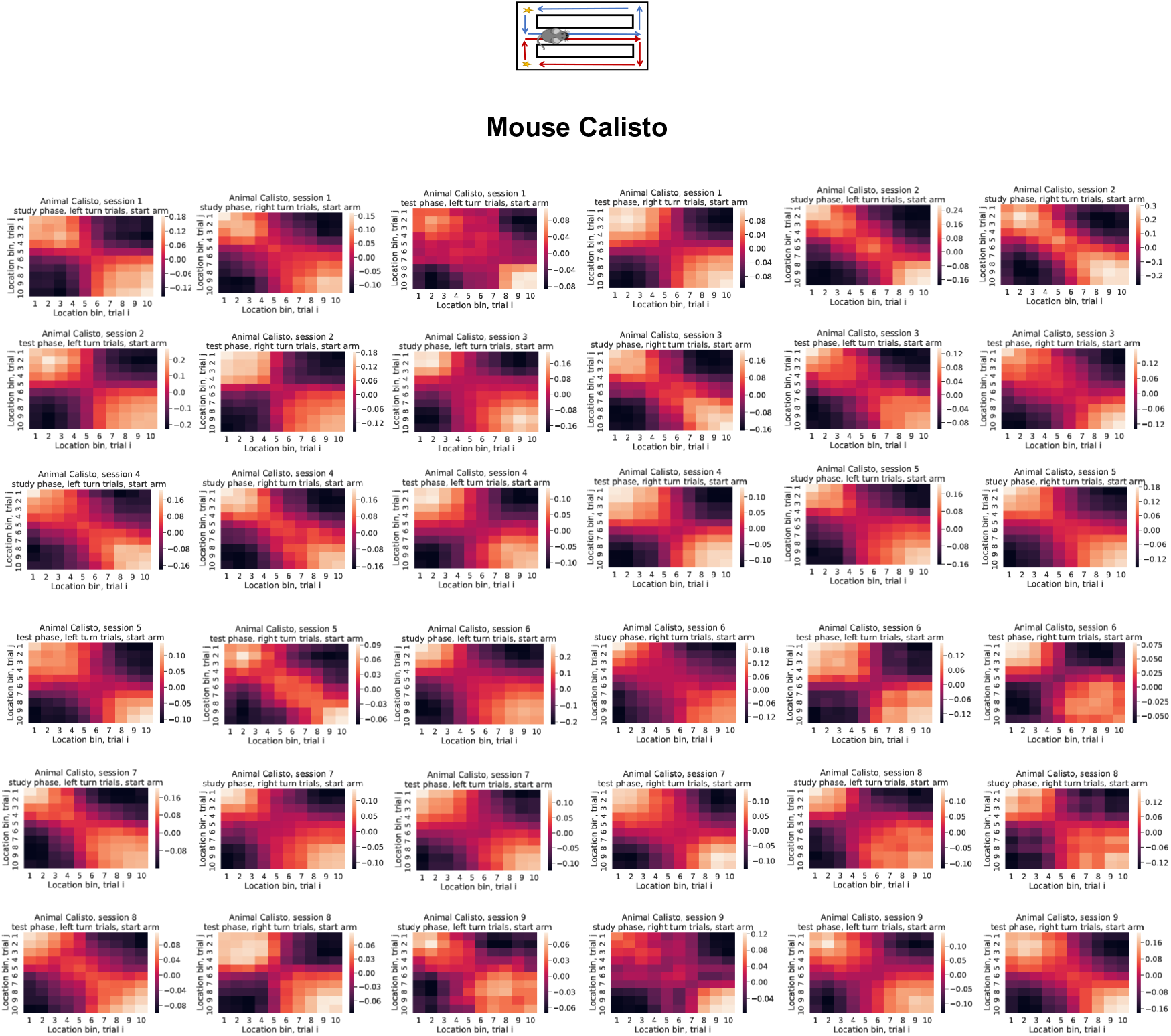
The cross-trial correlations for each individual session, task phase and turn direction in the spatial alternation task. Data for mouse Calisto.

**Figure S25.**
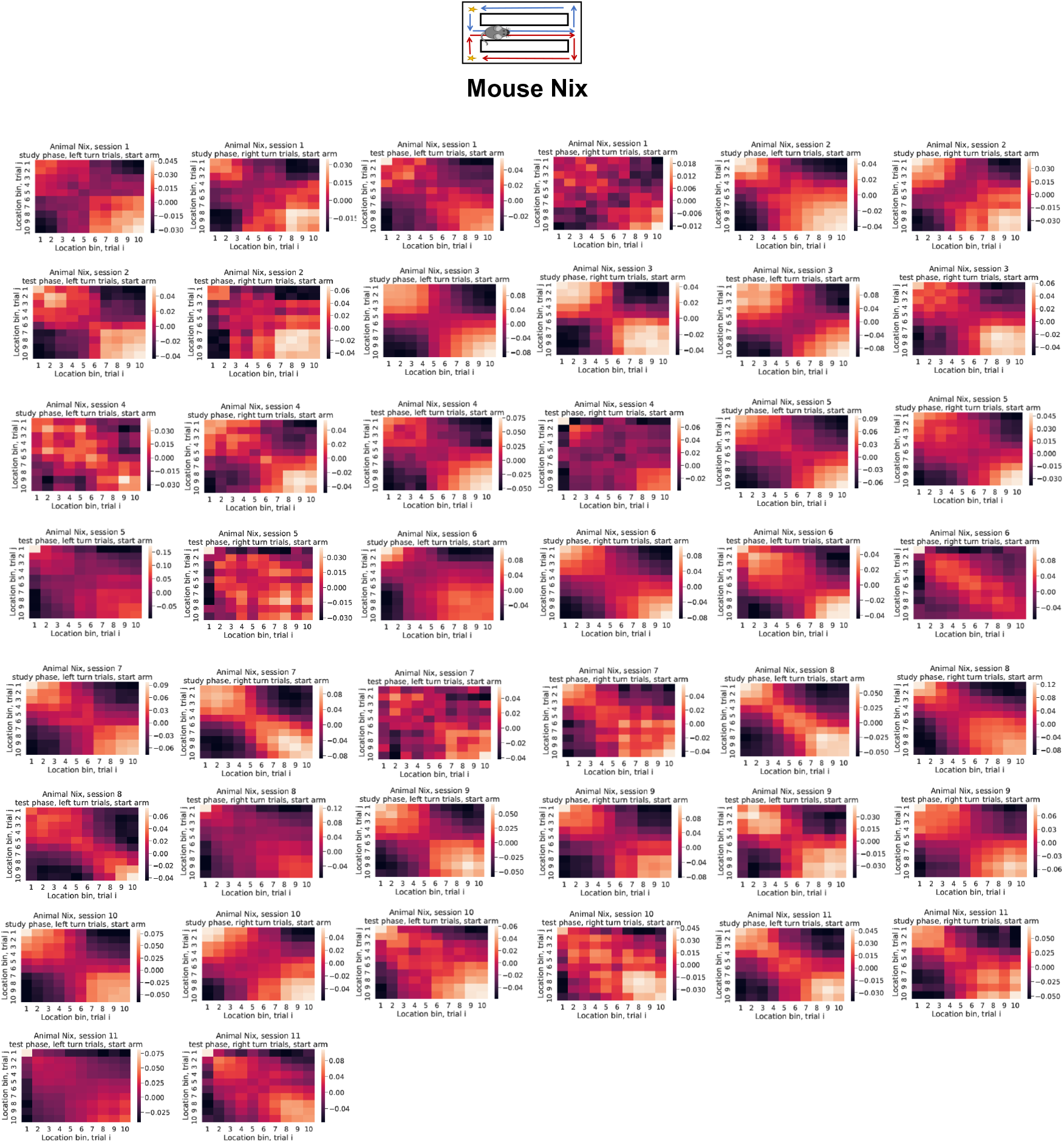
The cross-trial correlations for each individual session, task phase and turn direction in the spatial alternation task. Data for mouse Nix.

**Figure S26.**
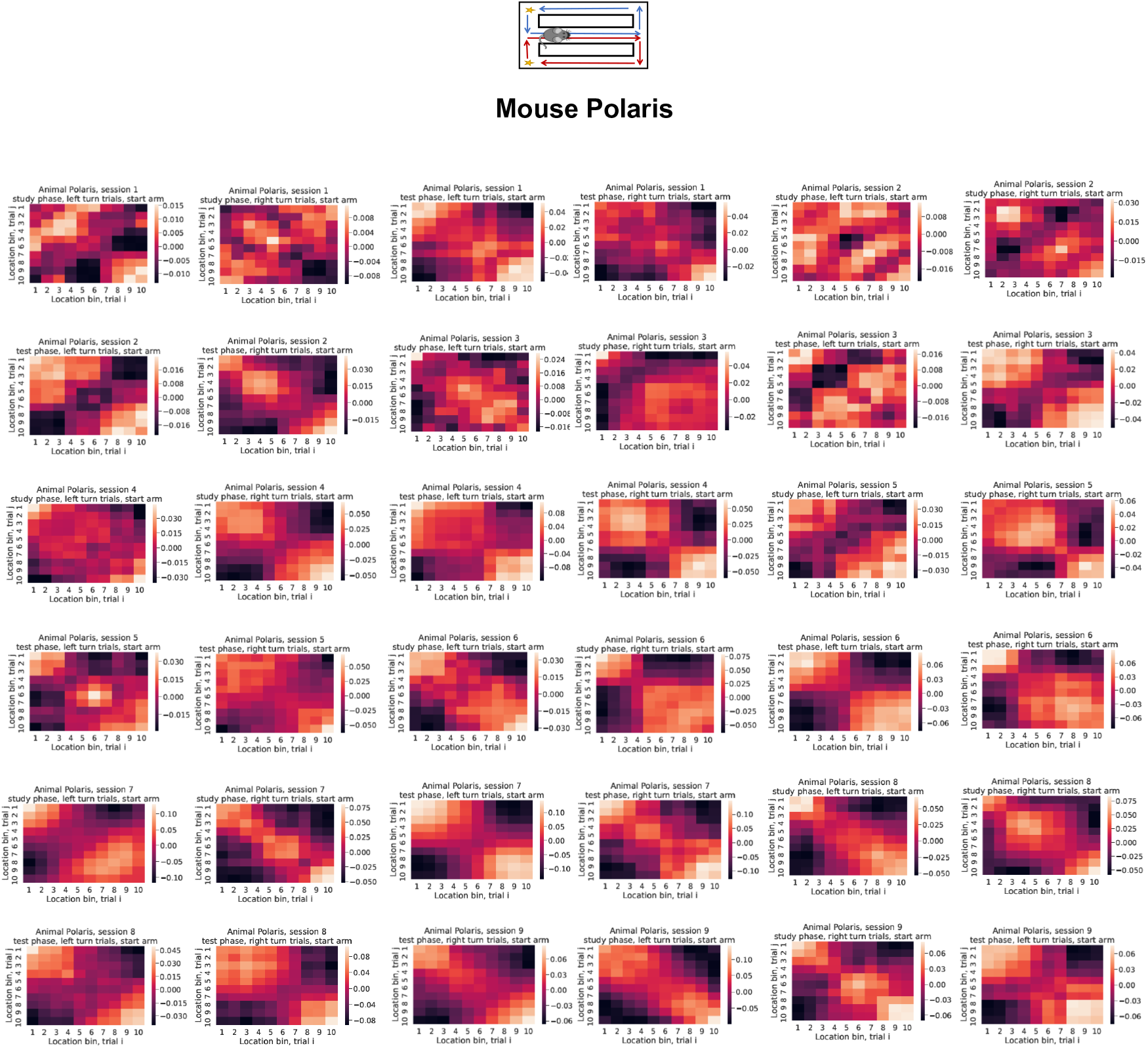
The cross-trial correlations for each individual session, task phase and turn direction in the spatial alternation task. Data for mouse Polaris.

**Figure S27.**
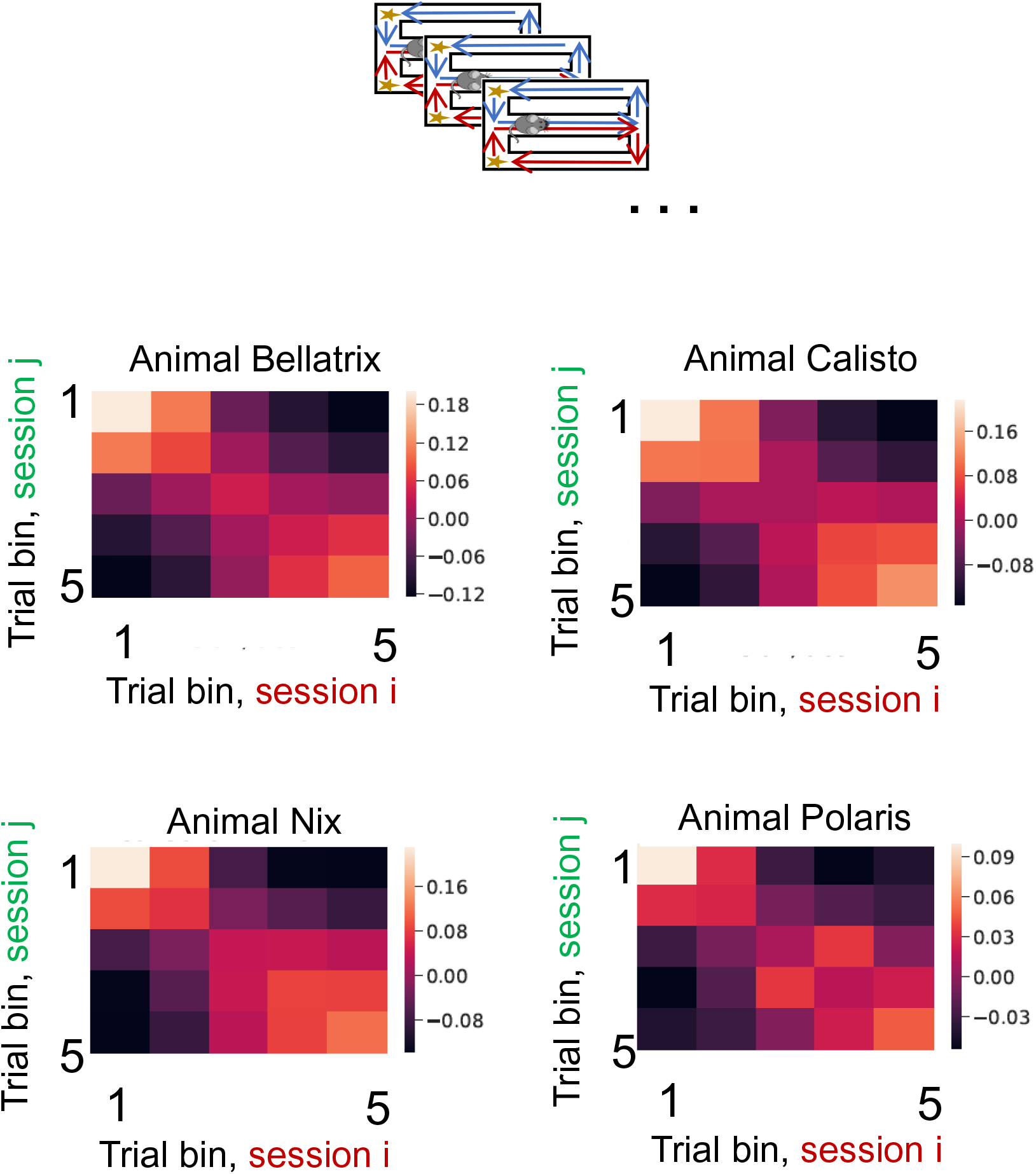
The cross-session correlations for each individual mouse in the spatial alternation task.

**Figure S28.**
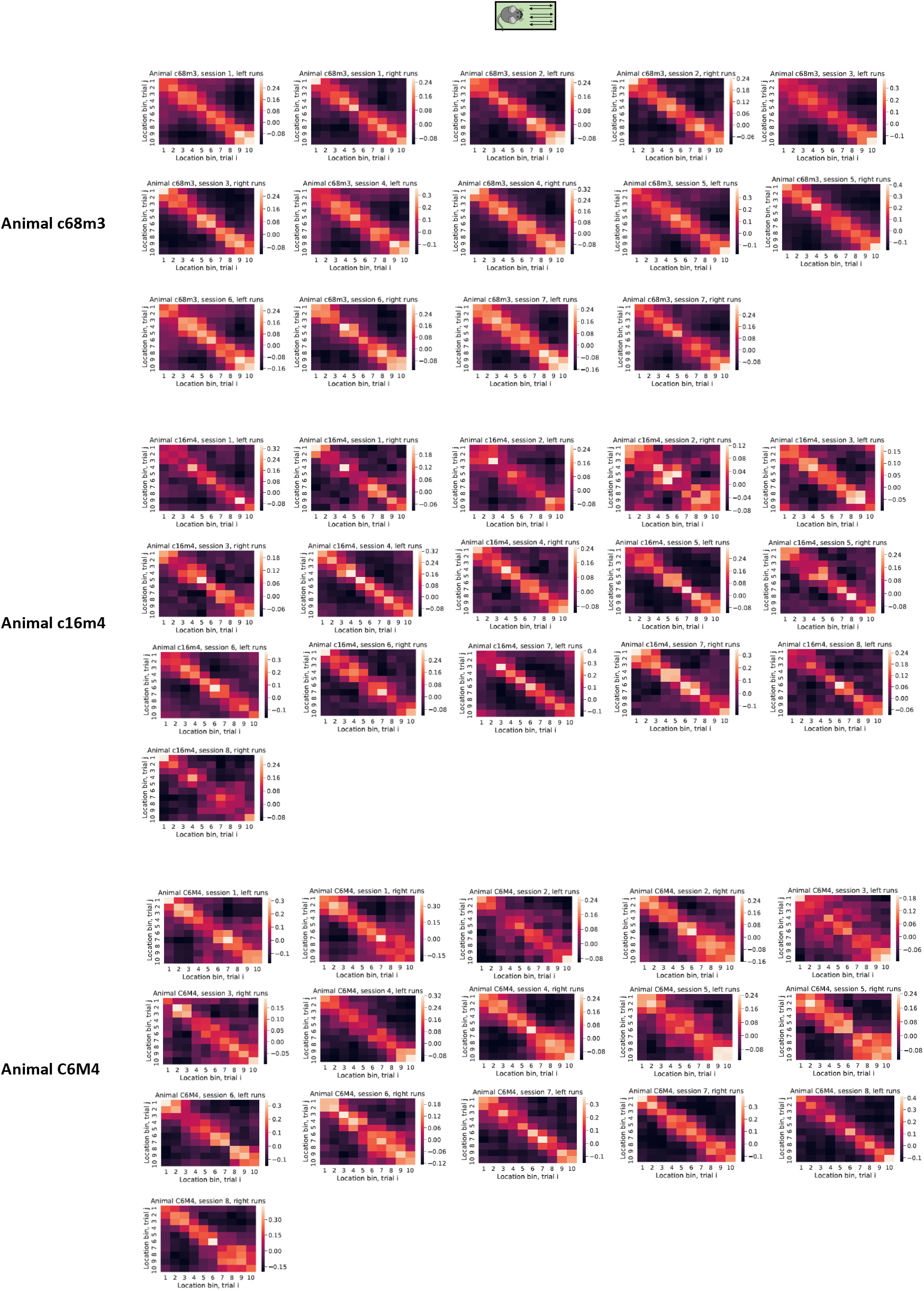
The cross-trial correlations for each individual session and running direction in the linear track task.

**Figure S29.**
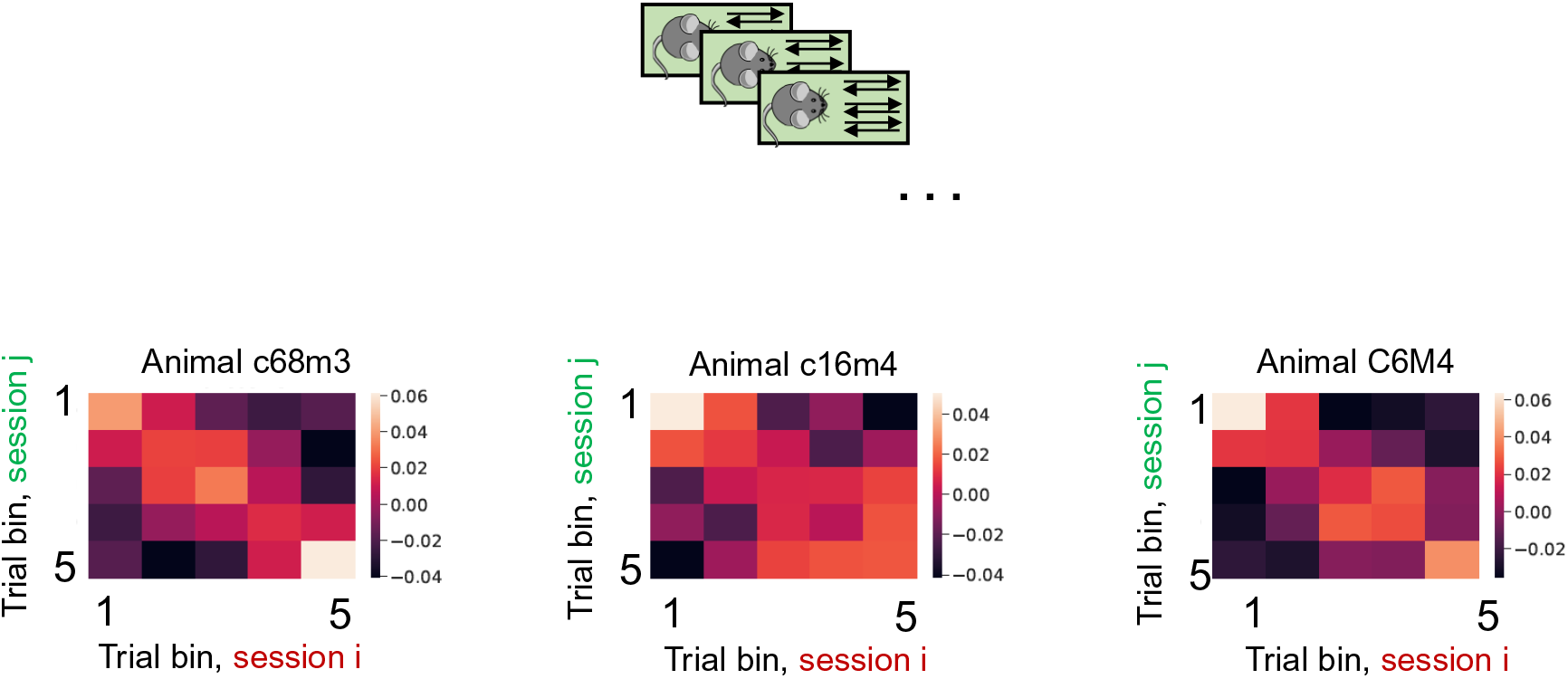
The cross-session correlations for each individual mouse in the linear track task.

